# Cadherin-11 is required for neural crest determination and survival

**DOI:** 10.1101/2020.05.18.066613

**Authors:** Subrajaa Manohar, Alberto Camacho, Crystal D. Rogers

## Abstract

Neural crest (NC) cells are multipotent embryonic cells that form melanocytes, craniofacial bone and cartilage, and the peripheral nervous system in vertebrates. NC cells express many cadherin proteins, which control their specification, epithelial to mesenchymal transition (EMT), migration, and mesenchymal to epithelial transition. Abnormal NC development leads to congenital defects including craniofacial clefts as well as NC-derived cancers. Here, we identify the role of the type II cadherin protein, Cadherin-11 (CDH11), in early chicken NC development. CDH11 is crucial for NC cell migration in amphibian embryos and is linked to cell survival, proliferation, and migration in cancer cells. It has been linked to the complex neurocristopathy disorder, Elsahy‐Waters Syndrome, in humans. Using immunohistochemistry (IHC), we determined that CDH11 protein has dynamic expression that is first co-localized with neural progenitors in early embryos and subsequently upregulated specifically in NC cells as they are specified in the dorsal neural tube prior to migration. We identified that loss of CDH11 led to a reduction of bonafide NC cells in the dorsal neural tube combined with defects in cell migration and survival. Loss of CDH11 increased p53-mediated programmed-cell death, and blocking the p53 pathway rescued the NC phenotype. Our findings demonstrate an early requirement for CDH11 in NC development, and may increase our understanding of early cadherin-related NC developmental defects.

**Summary:** Chicken Cadherin-11 (CDH11), which is expressed in neural crest (NC) cells prior to NC cell migration, is necessary for the determination and survival of the premigratory NC population.

## Introduction

Cadherin proteins are calcium dependent cell-cell adhesion molecules, which are essential for the development and maintenance of embryonic tissues (Giger and David, 2017; Taneyhill and Schiffmacher, 2017). Cadherins are single pass transmembrane proteins that contain a calcium-binding extracellular domain as well as a cytoplasmic domain which links with three catenin family proteins (α, β, and p120) and the actin cytoskeleton (Gul et al., 2017). These single pass transmembrane proteins have cytoplasmic regions that bind to β-catenin, p120 catenin (σ-catenin), and **α**-catenin, which directly links the actin cytoskeleton to the extracellular regions (Brasch et al., 2012). Classical cadherins are divided into types I and II. Type I cadherins include epithelial cadherin (CDH1) and neural cadherin (CDH2) among others, which have recently been implicated in both central nervous system (CNS) development in chick and zebrafish embryos (Dady et al., 2012; Miyamoto et al., 2015), and NC specification and EMT in frog, fish, and chicks (Huang et al., 2016; Piloto and Schilling, 2010; Rogers et al., 2013; Rogers et al., 2018). Type II cadherins include Cadherin 7 (CDH7), Cadherin 11 (Osteoblast-cadherin/CDH11) and Cadherin-6B (CDH6B), which have been linked to CNS patterning, NC cell delamination, EMT, and migration during embryonic development (Kashef et al., 2009; Liu et al., 2008; Schiffmacher et al., 2014; Schiffmacher et al., 2016; Taneyhill et al., 2007). With varied timing and onset of protein expression, the regulation and function of cadherin proteins is clearly important for normal development of the CNS, NC cells, and NC derivatives.

Here, we focus on the identifying the role of Cadherin-11 (CDH11) during early avian embryogenesis. CDH11 has been identified as a Wnt signaling target and effector in developmental and disease systems (Chalpe et al., 2010; Hadeball et al., 1998; Satriyo et al., 2019). Originally defined as a mesenchymal marker with no expression in the undifferentiated neural tube in mouse embryos (Hoffmann and Balling, 1995), the transcript was subsequently reported in developing mouse neuroepithelia (Kimura et al., 1995; Kimura et al.), as well as migratory NC cells in chick and frog embryos (Chalpe et al., 2010; Vallin et al., 1998). It has been identified as a major regulator of NC migration in *Xenopus* embryos (Abbruzzese et al., 2016; Kashef et al., 2009; Langhe et al., 2016; Vallin et al., 1998) and has been linked to tumor growth, cell survival, and the epithelial to mesenchymal transition (EMT) in disease models (Piao et al., 2016; Row et al., 2016; Yoshioka et al., 2015). Both reduced and increased levels of CDH11 are linked to patient survival and reduced metastasis in numerous cancers, however, its role is contrasting in different cancer cell types (Carmona et al., 2012; Lee et al., 2013). Specifically, high levels of CDH11 expression have been linked to poor prognosis in gastric cancer and triple-negative breast cancer (Chen et al., 2018; Satriyo et al., 2019), yet it maintains a pro-apoptotic tumor suppressor role in others (Li et al., 2012; Marchong et al., 2010). Although studies have linked CDH11 to NC migration in early development, there is less known about how the protein regulates NC induction, specification, maintenance, or survival.

NC cells are a vertebrate-specific population of stem-like cells that form craniofacial bone, cartilage, pigment cells, and the peripheral and enteric nervous systems (Hutchins et al., 2018; Rogers and Nie, 2018). In avian embryos, NC cells begin as tightly adherent neuroepithelial cells in the dorsal neural tube but detach from each other and the basal lamina to migrate using the process of EMT, which is controlled by alterations in the expression of type I and II cadherin proteins (Rogers et al., 2013; Scarpa et al., 2015; Taneyhill et al., 2007). Abnormal NC development can cause congenital defects known as neurocristopathies, which include cleft palate, craniofacial abnormalities, albinism, and defects in the enteric and peripheral nervous systems among others (Lopez et al., 2018; Reissmann and Ludwig, 2013). Bi-allelic mutations in CDH11 have specifically been linked to Elsahy-Waters syndrome, which is a combination of abnormal craniofacial developmental morphologies including those likely induced by neurocristopathies (Harms et al., 2018). The processes of NC specification and EMT are tightly regulated at multiple levels by signaling molecules (Bhattacharya et al., 2018), epigenetic modifiers (Hu et al., 2012), transcription factors (Simoes-Costa et al., 2015), and adhesion molecules (Abbruzzese et al., 2016; Rogers, 2018; Schiffmacher et al., 2016) to prevent developmental defects. Previous studies showed that perturbation of factors involved in this process can directly affect the formation and migratory ability of NC cells. Studies also identified links between cadherin proteins and NC cell migration and differentiation, however, there is little known about how type I or II cadherin proteins regulate premigratory NC development. Our study focuses on the role of CDH11 in NC formation (specification/determination), maintenance, and survival.

Here we examine the expression and function of CDH11 during early chicken development. Using IHC, we show the spatiotemporal localization of CDH11 in relation to pre-and post-migratory NC cells marked by PAX7 and HNK1. CDH11 is expressed in the neural tube prior to the onset of NC formation and is upregulated in the dorsal neural tube, co-localizing with PAX7 during NC specification (Hamburger Hamilton (HH) stage 8). As NC cells begin to undergo EMT (HH 9-10), CDH11 levels are increased in the premigratory and early migratory NC cells. Loss of CDH11 expression reduces the premigratory NC population marked by PAX7, SOX9, SNAI2, and SOX10, and increases membrane-associated CDH1, F-actin, and p53-mediated apoptosis in the presumptive NC regions. Our results indicate that the upregulation of CDH11 in the dorsal neural tube prior to NC migration is necessary for NC cell determination, migration, and cell survival.

## Results

### CDH11 is expressed during NC cell formation

CDH11 function has been extensively examined during NC migration and EMT in amphibian embryos (Abbruzzese et al., 2016; Borchers et al., 2001; Hadeball et al., 1998; Kashef et al., 2009; Koehler et al., 2013; McCusker et al., 2009; Vallin et al., 1998), but less is known about its endogenous expression and role in amniotes more specifically, developing avian embryos. To define the stage at which CDH11 proteins function in avian NC development, protein lysate was collected at multiple stages and we used western blot analysis to define the relevant stages. Protein from multiple stages of chicken embryos (HH4-6, HH8-10, and HH11-12) and from tailbud stage *Ambystoma mexicanum* (axolotl) was run to verify relative CDH11 protein size and stage of expression. We tested two antibodies against CDH11; a previously verified monoclonal mouse antibody against recombinant intracellular peptide of human CDH11 (Chalpe et al., 2010) and a rabbit polyclonal antibody directed against human CDH11 used previously in mouse tissue (Chang et al., 2017) (Fig. S1A). The mouse antibody identified two bands (potentially different isoforms) between the sizes of 110-135 kD while the rabbit antibody identified only the larger band and did not recognize axolotl CDH11. CDH11 protein appears to be expressed by stages HH4-6 and continues its expression through HH11-12 (Fig. S1A). Therefore, due to its specificity, we used the rabbit antibody to visualize CDH11 in our experiments. We also verified specificity of the antibody by performing CDH11 loss and gain of function experiments and visualizing reduction or exogenous expression of the protein (Fig. S2).

To characterize the spatiotemporal localization of CDH11 in early avian development, we performed IHC using anti-CDH11 in conjunction with previously characterized markers of NC cells (PAX7 and HNK1) (Basch et al., 2006; Del Barrio and Nieto, 2004). IHC revealed that CDH11 protein is expressed prior to NC specification. In 1 somite stage (SS) embryos, CDH11 is expressed in the neural plate (Fig. 1A, D, Fig. S1B-D’), but is markedly absent from the neural plate border, which expresses PAX7 (Fig. 1B, D). At this stage, CDH11 does overlap with the mesenchymal mesodermal cells marked by HNK1 (Fig. 1C, D). As the neural tube begins to fuse at 5 SS, CDH11 expression is maintained throughout the neural tube and co-localizes with a subset of PAX7-positive cells in the dorsal neural tube (Fig. 1E, F, H, Fig. S1E-F’). At this stage, CDH11 is discrete from HNK1-positive cells (Fig. 1G, H). At 7 SS, NC cells begin to delaminate and undergo EMT, CDH11 is upregulated in the most proximal PAX7-positive cells (Fig. 1I, J, L, Fig. S1G, G’). As HNK1 expression begins in the early migrating NC cells, the leading cells co-express CDH11 (Fig. 1K, L). In 9 SS embryos NC cells begin to migrate away from the midline, and all migratory cells are positive for both CDH11 and PAX7 (Fig. 1M, N, P), while the most lateral cells express HNK1 (Fig. 1O, P). Focusing on CDH11-postive cells at 9 SS shows that CDH11 is localized to the cell membranes as NC cells begin migrating from the midline (Fig. 1M’-P’). At 15 SS, CDH11 remains in the neural tube and the migratory NC cells, co-localizing with PAX7 (Fig. 1Q, R, T, Fig. S1H-I’), however, the cellular localization in later migratory NC appears more punctate and less membrane-bound (Fig. 1Q’-T’). Previous studies in *Xenopus* explants and transplanted NC cells showed that CDH11 must be cleaved extracellularly to allow for normal NC cell migration, and our results support those studies by demonstrating the endogenous intracellular expression changes in chick NC cells (Abbruzzese et al., 2016; Mathavan et al., 2017; McCusker et al., 2009). These data confirm that CDH11 is expressed during NC cell EMT and migration, but introduce novel expression in epithelial premigratory NC cells suggesting that CDH11 may play an earlier role in NC development.

**Figure 1.**
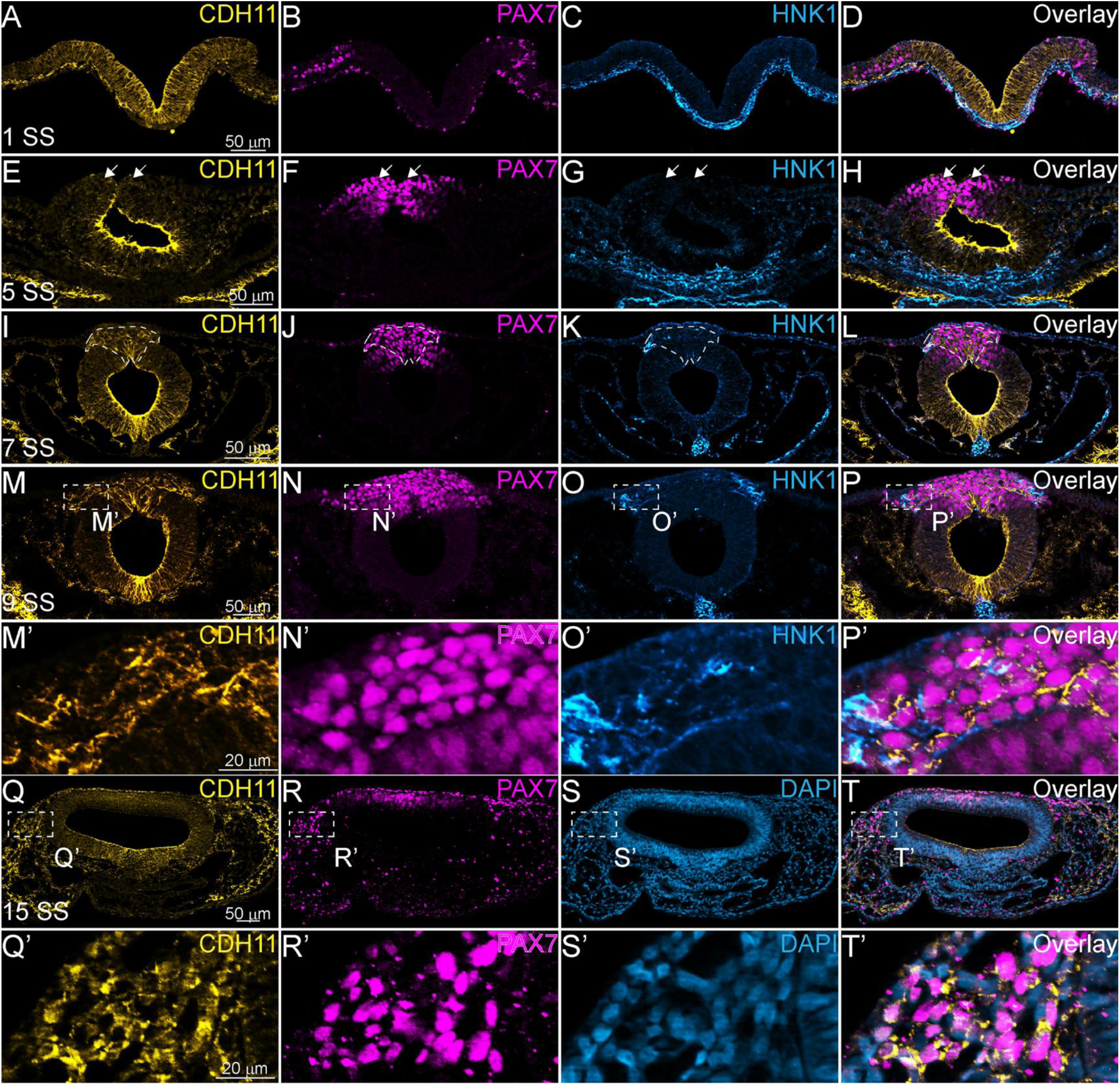
CDH11 is expressed in the neural tube and NC cells during multiple stages of chick development. To determine the stages at which CDH11 may function in NC development, Immunohistochemistry (IHC) was performed to visualize its endogenous expression in chick embryos. (A-T’) IHC using rabbit anti-CDH11 (yellow; A, E, I, M, M’, Q, Q’), PAX7 (pink; B, F, J, N, N’, R, R’), and HNK1 (blue; C, G, K, O, O’, S, S’) at (A-E) 1 somite stage (SS), (E-H) 5 SS, (I-L) 7 SS, (M-P’) 9 SS, and (Q-T’) 15 SS. (A-H) From 1SS to 5SS, CDH11 is localized throughout the neural plate/tube, but does not co-localize with either HNK1 in the mesoderm or PAX7 in the neural plate border at 1 SS. White arrows in E-H indicate lack of accumulation of CDH11 in the dorsal neural tube at this stage. (I-L) At 7 SS and (M-P) 9 SS CDH11 expression is upregulated and co-localized with PAX7 and HNK1 proteins in the premigratory and migratory NC cells in addition to its expression in the cranial mesenchyme. (M’-P’) Zoom in of dashed box in (M-P). (Q-T) At 15 SS CDH11 is weakly expressed in the neural tube and maintains expression in the migratory NC cells, overlapping with PAX7. (Q’-T’) Zoom in of dashed box in (Q-T). The early expression of CDH11 prior to migration suggests that it has a role in premigratory NC cell development. Scale bars in A-D as indicated in A, E-H in E, I-L in I, M-P in M, M’-P’ in M’, Q-T in Q, and Q’-T’ in Q’.

### CDH11 is necessary for NC cell population maintenance

The expression of CDH11 in premigratory NC cells suggests that it may play an earlier role in development than to drive migration. To understand the stage at which CDH11 is necessary for NC cell development, and to determine whether CDH11 was required for induction, specification, or determination, in addition to its role in migration, a time-course experiment was performed. We used a translation-blocking CDH11MO, which effectively reduced CDH11 protein on the injected side of the embryo compared to wildtype localization and the uninjected (UI) side (Fig. 2, Fig. S1A-C). CDH11 expression was inhibited at gastrula stage (HH 4), prior to the expression of the neural plate border marker, PAX7. After injection with CDH11MO, embryos were collected at stages HH5, and HH7 (3 somite stage (SS)) through HH10 (10 SS) and IHC for PAX7 was performed. At HH5 and 3 SS, PAX7 expression in the neural plate border and NC progenitors was unchanged even in the absence of CDH11 (Fig. 2A-B’, I). In these embryos, the UI and CDH11MO-injected sides had approximately the same number of cells (Fig. 2A-2B’, n= 14, p= 0.94). At HH8 (5 SS), the population of PAX7-positive NC cells was drastically reduced (Fig. 2C-2D’, n= 19, p= 0.0005). At HH10 (10 SS), there were less PAX7+ cells on the CDH11MO-injected side, suggesting that the NC cells did not recover prior to migration. (Fig. 2E-F, n= 7, p= 0.01). Compared to embryos injected with a non-specific control morpholino (Contmo), which had no change in PAX7 (Fig. 2G-H’, n= 14, p= 0.74), loss of CDH11 significantly reduced NC cells after induction, but during the stages at which NC specifier genes and proteins are normally upregulated. These data suggest that CDH11 is not required for NC induction, but that CDH11 is necessary for early NC development, specifically specification or determination, prior to its role in NC migration.

**Figure 2.**
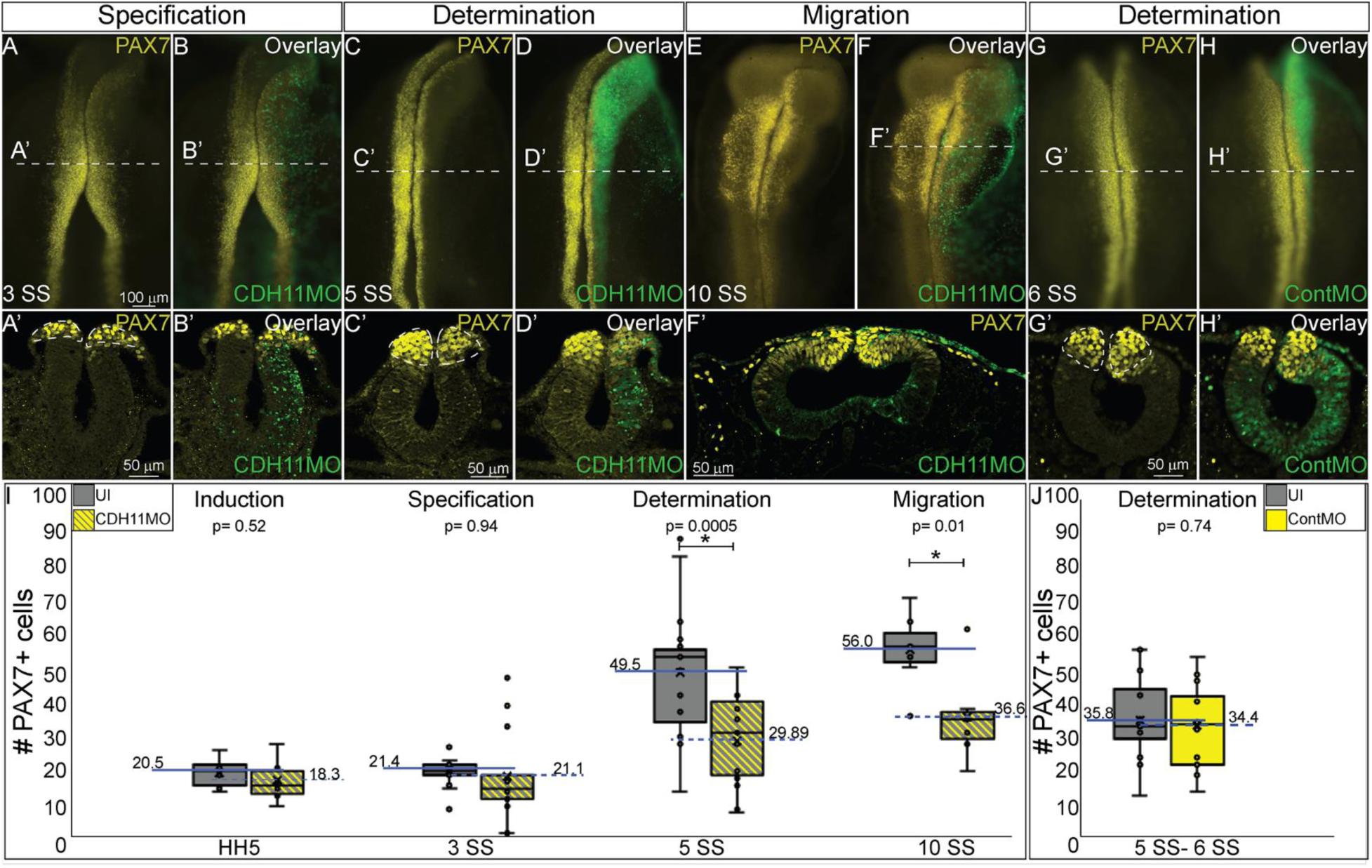
CDH11 is required for NC cell population determination. To determine the stage at which CDH11 is necessary for NC cell development, embryos were injected at HH4 and collected at multiple stages to analyze NC progenitor marker, PAX7. (A-F’) IHC for PAX7 on embryos injected unilaterally with CDH11MO at HH4 and collected between 3 SS and 10 SS. (G-H’) IHC for Pax7 in embryo injected unilaterally with ContMO and collected at 6 SS. (I, J) Actual PAX7+ cell counts of UI and morpholino-injected sides. (A) Whole mount IHC for PAX7 in HH8-(3 SS) embryo with (B) overlay with CDH11MO (green). (A’, B’) Transverse section of (A, B) with PAX7-positive NC cells circled. Mean number of cells at HH8-is 21.43 on UI and 21.14 on CDH11MO-injected side, p= 0.94, n= 14. (C) Whole mount IHC for PAX7 in HH8 embryo with (C) overlay with CDH11MO (green). (C’, D’) Transverse section of (C, D) with PAX7-positive NC cells circled. Mean number of cells at HH8 is 49.53 on UI and 29.89 on CDH11MO-injected side, p= 0.0005, n= 19. (E) Whole mount IHC for PAX7 in HH10 embryo with (F) overlay with CDH11MO (green). (F’) Transverse section of (F) with PAX7-positive NC cells. Mean number of cells at HH10 is 56.00 on UI and 36.57 on CDH11MO-injected side, p= 0.01, n= 7. (G) Whole mount IHC for PAX7 in HH8 embryo with (H) overlay with ContMO (green). (G’, H’) Transverse sections of (G, H) with PAX7-positive NC cells circled. Mean number of cells on UI side is 35.8 and on ContMO-injected side is 34.4, p= 0.74, n= 14. Dashed circles indicate area of UI side and are mirrored on injected sides to demonstrate changes in the NC cell population density. At 5 SS the NC cell population is less dense in the CDH11MO-side compared to UI when compared to 3 SS and ContMO-injected embryos. Scale bars are as marked (100 µm for whole mount and 50 µm for sections). Anterior to top in all whole mount images, dorsal to top in all sections. Loss of CDH11 reduces the PAX7-positive NC cell population after induction (I, HH5-3SS) and at a point between specification and determination (5 SS).

### Loss of CDH11 reduces the definitive NC cell population

At the stages tested, CDH11 is expressed in both the neural tube and definitive NC cells. First, to determine if the CDH11-knockdown phenotype was NC-specific or if loss of CDH11 affected all ectodermal derivatives, we analyzed the expression of SOX2, a neural tube progenitor marker, after CDH11 knockdown (Fig. 3A-C). Expression of SOX2-positive cells was unaffected in the CDH11MO-innjected side compared to the UI-side (Fig. 3A-C, n= 5, p= 0.9). As reported by multiple groups, in the NC gene regulatory network (GRN) the factors in the neural plate border (i.e. PAX7) drive the expression of bonafide NC markers (SNAI2, SOX9, SOX10, etc.) as neurulation proceeds (Williams et al., 2019). These factors are then responsible for altering the expression of specific cadherin proteins and allowing for NC cell EMT and migration (Taneyhill et al., 2007; Taneyhill and Schiffmacher, 2017). To determine if loss of CDH11 universally reduced the NC cell population by reducing NC progenitors and definitive NC cells, embryos were unilaterally injected with CDH11MO, and IHC for markers of bonafide NC cells (SOX9, SNAI2, SOX10) was performed at HH8-9 (5 SS to 7 SS). Loss of CDH11 significantly reduced SOX9-postive NC cells by 41% (Fig. 2J-L, n= 11, p= 0.04). SNAI2-positive cells were reduced by 34% (Fig. 2M-O, n= 16, p= 0.001). SOX10-positive NC cells were reduced by 41.7% (Fig. 2P-R, n= 19, p= 0.01). The loss of CDH11 reduces both progenitors and definitive NC cells prior to NC cell migration. However, each NC specifier protein drives specific programs with regards to NC cell development. SNAI2 inhibits CDH6B and CDH1 expression to drive cell migration (Taneyhill et al., 2007; Tien et al., 2015) and it is also linked to the inhibition of apoptotic activity in NC cells (Tribulo et al., 2004). While the SOXE proteins, SOX9 and SOX10 are linked with the progression of NC cell migration (Cheung and Briscoe, 2003). To determine the mechanisms downstream of CDH11 knockdown that led to a reduction in the NC population, we next analyzed cell death and proliferation.

**Figure 3.**
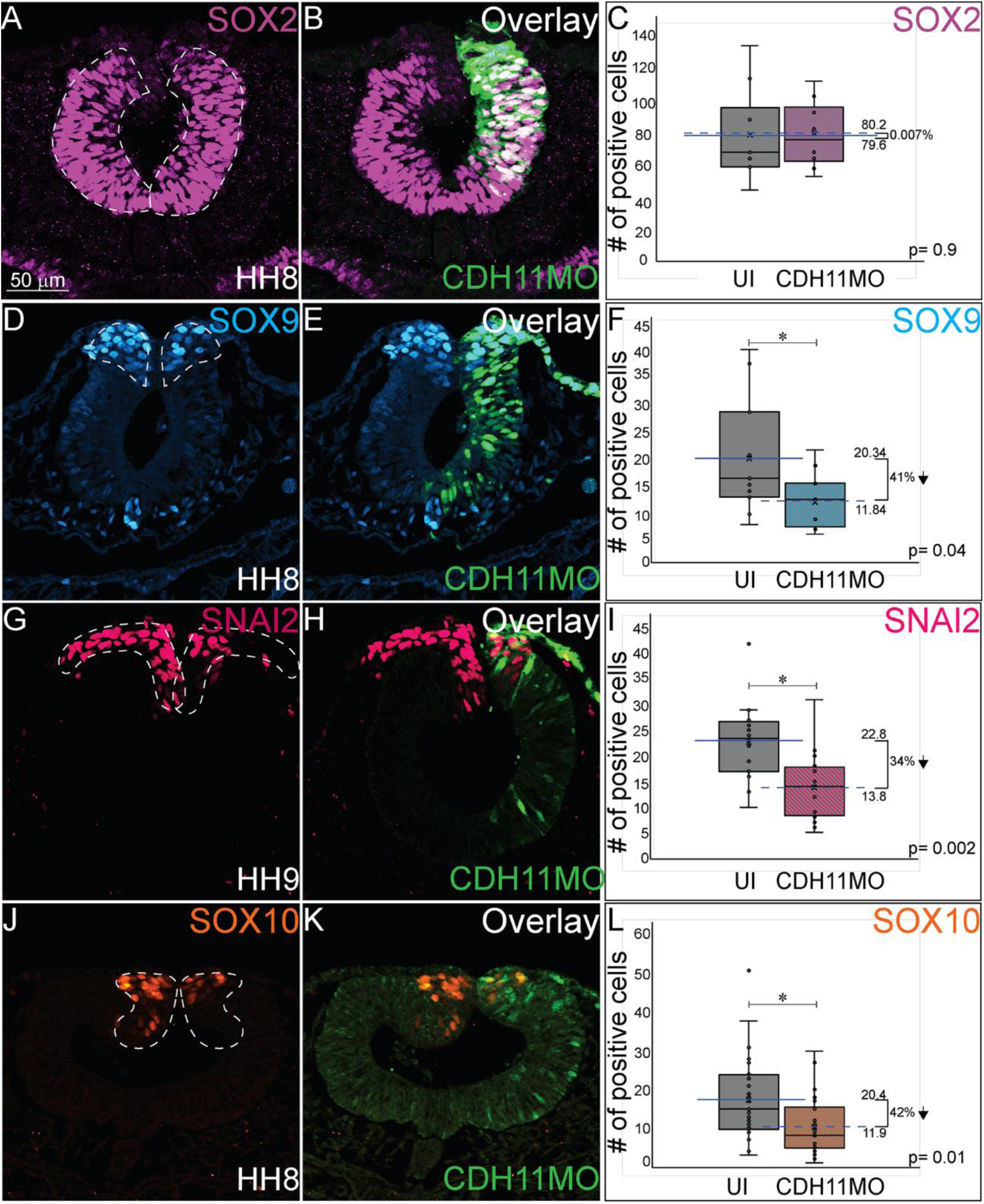
Loss of CDH11 reduces definitive NC cells. To determine if loss of CDH11 affected neural progenitors and definitive NC cells in addition to NC progenitors, embryos were injected unilaterally with CDH11MO or ContMO and IHC was performed for neural progenitors (SOX2) and definitive NC cells (SOX9, SNAI2, SOX10). (A) IHC for SOX2 in transverse section from HH8 embryo with (D) overlay with CDH11MO (green). (C) Graph showing difference between UI and CDH11MO-injected sides. Mean number of SOX2+ cells is 80.20 on UI and 79.50 on CDH11MO-injected side, p= 0.90, n= 5. (D) IHC for SOX9 in transverse section from HH8 embryo with (E) overlay with CDH11MO (green). (F) Graph showing difference between UI and CDH11MO-injected sides. Mean number of SOX9+ cells is 20.34 on UI and 11.85 on CDH11MO-injected side, p= 0.04, n= 11. (G) IHC for SNAI2 in transverse section from HH9 embryo with (H) overlay with CDH11MO (green). (I) Graph showing difference between UI and CDH11MO-injected sides. Mean number of SNAI2+ cells is 22.75 on UI and 13.75 on CDH11MO-injected side, p= 0.001, n= 16. (J) IHC for SOX10 in transverse section from HH8 embryo with (K) overlay with CDH11MO (green). (L) Graph showing difference between UI and CDH11MO-injected sides. Mean number of SOX10+ cells is 20.35 on UI and 11.87 on CDH11MO-injected side, p= 0.01, n= 18. Reducing CDH11 significantly reduces the entire population of premigratory NC cells without affecting the SOX2-positive neural tube progenitors. All graphs show mean (indicated on graph) and median (line within graph). Scale bar for A, B, D, E, G, H, J, K indicated in A.

### CDH11 is required for NC cell survival

The reduction in PAX7-positive NC cells after CDH11 knockdown could be caused by two cellular responses. Our results show that CDH11 reduction leads to a loss of both NC progenitors and definitive NC cell markers. To determine if this phenotype was a result of increased cell death (or reduction of cell survival) or of a reduction NC cell proliferation (or reduced population growth), we unilaterally injected CDH11MO or ContMO into HH4 chicken embryos and performed IHC for markers of cell death and cell proliferation. To determine the requirement for CDH11 in NC cell survival, IHC was performed for activated Caspase-3 (*Casp3), a protein that initiates programmed cell death downstream of p53 (Zou et al., 1999). Total numbers of *Casp3-positive apoptotic bodies was counted in the injected and uninjected sides of each embryo. Uninjected sides were normalized to 1 and the average fold increase was calculated. Expression of *Casp3 in apoptotic bodies was increased to 176.5% in CDH11MO-injected sides compared with the UI side (Fig. 4A-C, n=14, p= 0.04). Embryos injected with ContMO did not have a significant difference in *Casp3 expression or apoptotic bodies on the injected versus UI side (Fig. 4D-F, n= 14, p= 0.97). Loss of CDH11 is correlated with an increase in cell death on the injected side of the embryo, and therefore the presence of CDH11 may be necessary for NC cell survival.

**Figure 4.**
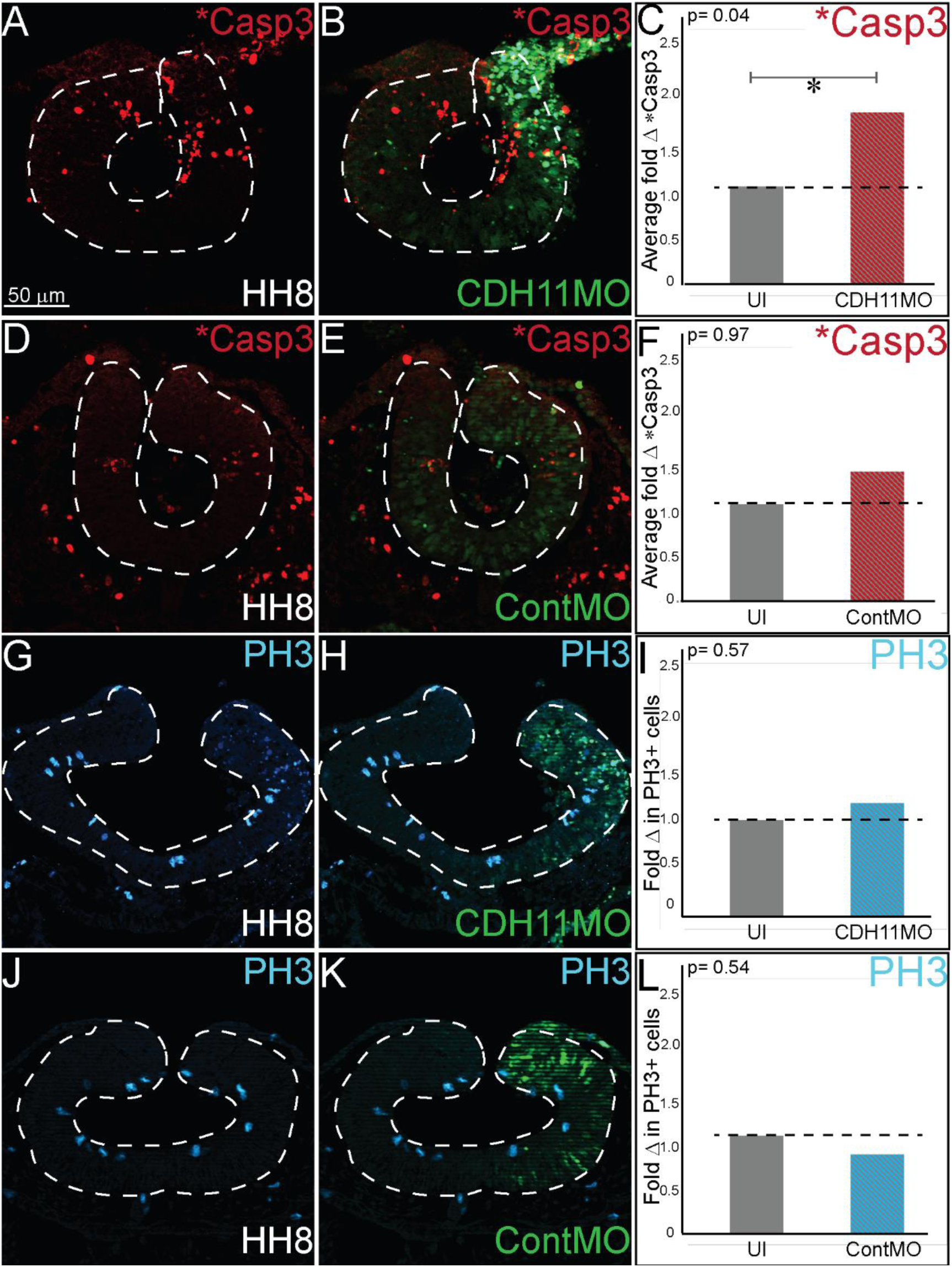
Loss of CDH11 increases cell death. To determine the cause of reduced NC cell population after CDH11 knockdown, embryos were injected unilaterally with CDH11MO or ContMO and IHC was performed for activated caspase 3 (*Casp3) to mark apoptotic cells or phosphorylated histone H3 (PH3) to mark mitotic cells. (A) IHC for *Casp3 in transverse section from HH8 embryo with (B) overlay with CDH11MO (green). (C) Graph showing fold difference between UI and CDH11MO-injected sides. Mean number of *Casp3+ apoptotic bodies is 17.90 on UI and 31.86 on CDH11MO-injected side and there is a 77% increase in *Casp3 expression on CDH11MO-injected side, p= 0.04, n= 14. (D) IHC for *Casp3 in transverse section from HH8 embryo with (E) overlay with ContMO (green). (F) Graph showing difference between UI and ContMO-injected sides. Mean number of *Casp3+ apoptotic bodies is 9.36 on UI and 9.36 on ContMO-injected side, p=0.97, n= 14. (G) IHC for PH3 in transverse section from HH8 embryo with (H) overlay with CDH11MO (green). (I) Graph showing difference between UI and CDH11MO-injected sides. Mean number of PH3+ cells is 6.20 on UI and 5.20 on CDH11MO-injected side, p= 0.50, n= 20. (J) IHC for PH3 in transverse section from HH8 embryo with (K) overlay with ContMO (green). (L) Graph showing difference between UI and ContMO-injected sides. Mean number of PH3+ cells is 5.88 on UI and 4.75 on ContMO-injected side, p= 0.54, n= 8. Loss of CDH11 increases cell death on injected side. All graphs show mean (indicated on graph) and median (line within graph). Scale bar for A, B, D, E, G, H, J, K indicated in A.

Previous results in *Xenopus* embryos linked loss of CDH11 to increased Wnt-dependent cell cycling and suggested that loss of CDH11 was positively correlated with NC cell proliferation (Koehler et al., 2013). To determine if NC cell proliferation was changed after CDH11 knockdown, we injected CDH11MO or ContMO unilaterally in HH4 embryos, and performed IHC for phosphorylated histone H3 (PH3), a marker of mitotic cells (Fig. 4G-L). We determined that neither CDH11MO (Fig. 4G-I, n= 20, p= 0.57) nor ContMO (Fig. 4J-L, n= 8, p= 0.54) significantly changed the number of PH3-positive cells suggesting that there is no consistent role for CDH11 in NC proliferation. The lack of discernible reduction in SOX2+ neuroepithelial cells after CDH11MO-injection (Fig. 3A-C) and significant reduction in the number of definitive NC cells (Fig. 3D-L) suggests that SOX2+ neuroepithelial cells are able to recover from the loss of CDH11 while NC cells are not. These data demonstrate a requirement for CDH11 in NC cell survival.

### Blocking p53-mediated apoptosis rescues NC cells

Previous studies identified SNAI2 and SOX9 as factors that prevent apoptosis in NC cells in *Xenopus* and chick embryos, respectively (Cheung et al., 2005; Tribulo et al., 2004). Due to the reduction in the expression of these proteins and subsequent increase in *Casp3 expression after CDH11 knockdown we hypothesized that blocking the p53-mediated apoptotic pathway would rescue the phenotype caused by reduction of CDH11. To confirm that the NC phenotype was specific to changes in CDH11 expression, we first compared embryos injected with CDH11MO alone to those co-injected with CDH11MO and full length CDH11-GFP and performed IHC for PAX7 expression (Fig. 5A, B, E, F, and K). Whereas loss of CDH11 reduced PAX7-expressing cells at stages after HH8 significantly (Fig. 5A, K, n= 18, p= 0.02), PAX7 expression was rescued by co-injection of CDH11-GFP (Fig. 5B, E, F, K, n= 13, p= 0.15). We next co-injected CDH11MO with a p53 translation-blocking morpholino (p53MO) to rescue the loss of PAX7. Injection of p53MO alone had little effect on PAX7 expression (Fig. 5C), but co-injection of CDH11MO with p53MO rescued PAX7 expression (Fig. 5D, G, H, K, n= 16, p= 0.48). To confirm that the renewed PAX7 expression was caused by a reduction in CDH11-p53-mediated cell death, we analyzed *Casp3 expression in CDH11MO and p53MO co-injected embryos. Whereas CDH11MO increased *Casp3 expression (Fig. 4A-C, Fig. 5L, n= 14, p= 0.04), injection of p53MO alone did not significantly change levels of *Casp3 in embryos (Fig. 5L, n= 10, p= 0.71). However, blocking p53 rescued the increased *Casp3 expression previously induced by the loss of CDH11 and reduced the number of apoptotic bodies so that there was no significant difference between the injected and UI sides (Fig. 5I, J, L, n= 12, p= 0.68). These data demonstrate that loss of CDH11 reduces NC cells due to increased p53-mediated cell death, and that blocking p53-mediated apoptosis can rescue the loss of NC cells.

**Figure 5.**
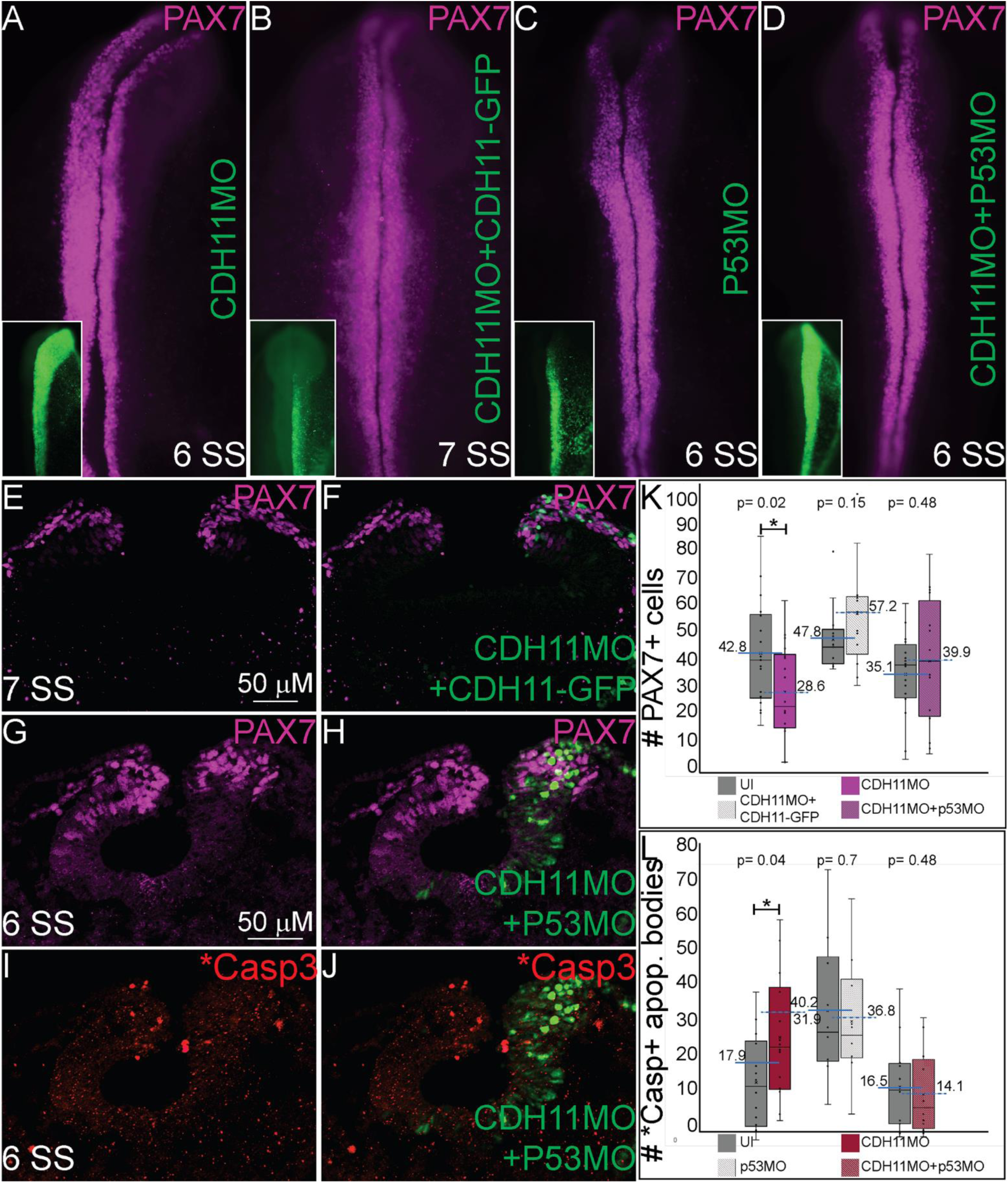
Blocking p53-mediated apoptosis rescues the NC fate. To determine if the NC and *Casp3 phenotypes are resulted from p53-mediated apoptosis, embryos were injected with multiple combinations of treatments to attempt to rescue the phenotype. (A-D) Whole mount IHC for PAX7 in HH8-HH9 embryos after (A) CDH11MO, (B) CDH11MO+CDH11-GFP, (C) p53MO, or (D) CDH11MO+p53MO. Inset shows CDH11MO injection (green). (E) IHC for PAX7 in transverse section from HH9 embryo with (F) overlay with CDH11MO+CDH11-GFP (green). (G) IHC for PAX7 in transverse section from HH9 embryo with (H) overlay with CDH11MO+p53MO (green). (I) IHC for *Casp3 in transverse section from HH8 embryo with (J) overlay with CDH11MO+p53MO (green). (K) Graph showing difference in PAX7 expression between UI and injected sides. Mean number of PAX7+ cells is 42.8 on UI and 28.6 on CDH11MO-injected side, p= 0.02, n= 18. Mean number of PAX7+ cells is 47.77 on UI and 57.23 on CDH11MO+CDH11-GFP-injected side, p= 0.15, n= 13. Mean number of PAX7+ cells is 35.06 on UI and 39.94 on CDH11MO+p53MO-injected side, p= 0.48, n= 16. (L) Graph showing difference in *Casp3 expression between UI and injected sides. Mean number of *Casp3+ cells is 17.90 on UI and 31.86 on CDH11MO-injected side, p= 0.04, n= 14. Mean number of *Casp3 + cells is 40.20 on UI and 36.80 on CDH11MO+CDH11-GFP-injected side, p= 0.71, n= 10. Mean number of *Casp3 + cells is 16.50 on UI and 14.17 on CDH11MO+p53MO-injected side, p= 0.68, n= 12. All graphs show mean (indicated on graph) and median (line within graph). Phenotypes were rescued by co-injection with full length CDH11 as well as by blocking p53 translation suggesting that the NC phenotype is due to cell death after loss of CDH11. Scale bars for E, f are as marked in E and G, H, I, and J are marked in G.

### CDH11 is required for normal NC cell morphology and migration

To investigate whether the early loss of CDH11 affected NC cell morphology *in vivo*, we analyzed the expression of CDH1, a membrane-bound type I cadherin protein that has been linked to NC cell specification (Rogers, 2018) and in NC cell EMT and migration in both frog and chick embryos (Huang et al., 2016; Rogers et al., 2013). CDH11MO was injected unilaterally, and cell morphology, CDH1 fluorescence intensity, and cell migration distance was measured in HH10 (10 SS) embryos. Loss of CDH11 enhanced the fluorescence of CDH1 in the dorsal neural tube on the injected side, and the CDH1/SOX9-positive cells appeared smaller and rounder than those on the UI side (Fig. 6A-A’’’, B, B’, I, n= 11, p= 0.02). We next measured the distance from the midline that PAX7 and SOX9-positive cells traveled in CDH11MO-injected cells compared to UI sides. CDH11MO-injected SOX9-positive cells migrated 31% less than the UI side (Fig. 6C, D, E, n= 17, p= 0.026) and PAX-positive cells migrated 43.8% less than the UI side (Fig. 6J, n= 19, p= 0.029). Our results suggest that CDH11 is necessary for NC specification or determination and survival prior to EMT. As a result of the early phenotype, cell morphology and migration remain affected at migratory stages.

**Figure 6.**
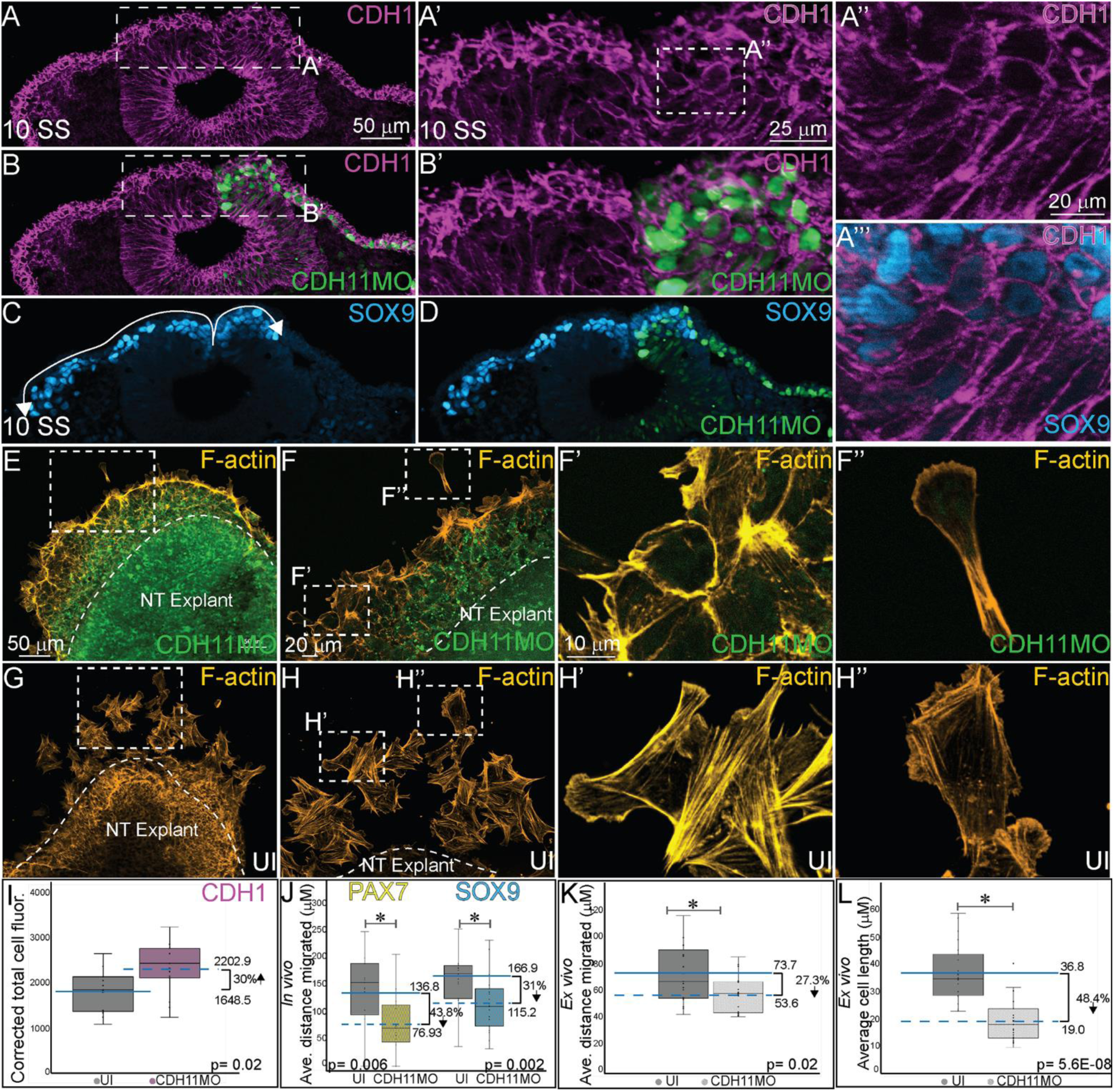
CDH11 knockdown affects cell morphology and NC cell migration. To determine if the NC determination and cell death phenotype affects cell morphology *in vivo* (A-D) embryos were injected unilaterally with CDH11MO and IHC was performed for E-cadherin (CDH1) to mark epithelial cells and SOX9 to mark definitive NC cells. To determine if the CDH11 knockdown phenotype affects cell morphology and migration *ex vivo* (E-H’’) Embryos were injected unilaterally with CDH11MO and the injected and UI sides were dissected from HH8 embryos, cultured on fibronectin coated slides, and stained for filamentous actin (F-actin). (A) IHC for CDH1 in transverse section from 10 SS embryo with (B) overlay with CDH11MO (green). (A’) Zoom in of dashed box from (A) showing rounded cell morphology in dorsal neural tube in CDH11MO-injected versus UI side. (B’) Overlay with CDH11MO. (A’’) Zoom in of dashed box from (A’) and (A’’’) overlay with SOX9-positive cells. (C) IHC for SOX9 in transverse section from the same 10 SS embryo from (A) with (D) overlay with CDH11MO (green) demonstrating reduced migration on CDH11-injected side. (E-F’’) Staining for F-actin in explant from embryo unilaterally injected with CDH11MO at (E) 20X and (F) 40X magnification. (F’) Zoom in of single follower cell from CDH11MO-injected explant. (F’’) Zoom in of single leading cell from CDH11MO-injected explant. Both cells are significantly closer to epithelial explant and smaller than UI cells. (G-H’’) Staining for F-actin in explant from UI side at (G) 20X and (H) 40X magnification. (H’) Zoom in of grouped follower cells from UI explant. (H’’) Zoom in of single leading cell from UI explant. (I) Graph showing *in vivo* difference in CDH1 between UI and CDH11MO-injected sides. Corrected mean total cell fluorescence of CDH1 in the dorsal neural tube is 1645.5 on UI and 2202.9 on CDH11MO-injected side, p= 0.02, n= 11. (J) Graph showing difference in migration *in vivo* from midline of PAX7 and SOX9-positive cells between UI and CDH11MO-injected sides. Average distance migrated away from midline by PAX7+ cells is 136.8 µm on UI and 76.93 µm on CDH11MO-injected side, p= 0.006, n= 11 cells. Average distance migrated away from midline by SOX9+ cells is 166.87 µm on UI and 115.22 µm on CDH11MO-injected side, p= 0.002, n= 19. (K) Graph showing average distance migrated *ex vivo* by cells from explant is 73.7 µm from UI explant and 53.6 µm from CDH11MO-injected explant, p= 0.02, n= 17 cells. (L) Graph showing average cell length is 36.8 µm in UI explants and 19.0 µm in CDH11MO-injected explants, p= 5.6E-08, n= 23 cells. Overall, loss of CDH11 significantly reduces NC cell population, affects their morphology, and reduces cell migration as a result. All graphs show mean (indicated on graph) and median (line within graph). Scale bars for A, B, C and D are as marked in A, A’, B’ are marked in A’, A’’ and A’’’ are marked in A’’, E and G are marked in E, F and H are marked in F, F’-H’’ are marked in F’.

We next performed NC explant assays to assess the morphology and migratory ability of the cells lacking CDH11 *ex vivo* to determine if the NC migration defects in embryos are due intrinsic or extrinsic properties. Embryos were injected unilaterally with CDH11MO at HH4. Neural tube explants were dissected from the embryos between HH8 (3 SS) and HH9 (6 SS), cultured for 8 hours, and then stained for filamentous actin (F-actin). Explants were repeated in triplicate and both leading edge and follower cells were measured (distance and size) individually. Cells lacking CDH11 migrated away from the explant 27.3% less than UI cells (Fig. 6E-F’’, K, n= 17, p= 0.02). The average migration distance from the epithelial explant in CDH11MO-inejcted cells was 53.6 µm while UI cells demonstrated a normal migratory ability, formed both lamellipodia and filopodia, and migrated approximately 73.7 µm from the explant (Fig. 6G-H’’, K). Very few cells were physically able to detach from the collective group in CDH11MO-injected explants, and most remained strongly adherent to the epithelial explant (compare Fig. 6E, F’ to 6F, F’’). In addition to exhibiting migration defects, cells lacking CDH11 were significantly smaller more rounded, lacked filopodia similar to previously described in *Xenopus* NC cells lacking CDH11 (Kashef et al., 2009), and were generally unable to detach from the neural tube explant (compare Fig. 6F’, F’’ to 6H’, H’’, n= 23, p= 5.6E-08). Loss of CDH11 causes p53-mediated cell death and reduces the number of bonafide NC cells in addition to causing morphological and migratory defects both *in vivo* and *ex vivo*. Our analyses add new information to previous discoveries, demonstrating that CDH11 is not solely required for migration, but plays an important role prior to NC cell emigration from the neural tube.

## Discussion

Defining the specific roles of cell adhesion proteins in early development is necessary as abnormal expression of these proteins is linked to cellular anomalies and congenital defects. With dynamic expression profiles, differential downstream signaling, and implications in cell survival, specification, and migration, cadherin proteins are involved in multiple aspects of embryonic development. Here, we examined the localization and function of CDH11 protein in early avian NC cells. CDH11 is expressed in the neural tube prior to NC cell formation, is upregulated in premigratory NC cells, and is maintained in migratory NC cells. Loss of CDH11 reduced both NC progenitor and definitive NC cell populations, specifically, cells that normally drive NC cell migration (SNAI2, SOX9, SOX10) and NC cell survival (SNAI2, SOX9). In contrast to previous studies, loss of CDH11 had little effect on cell proliferation but increased p53-mediated cell death in the neural tube, potentially as a result of losing SNAI2 and SOX9, and the increase in CDH1, preventing normal NC progression and migration. Inhibition of p53 in cells deficient for CDH11 rescued the loss of NC cells and the increase in *Casp3 (Fig. 7A). Similar to previous studies in frog embryos (Kashef et al., 2009; Langhe et al., 2016), CDH11-deficient cells exhibited migratory and morphological defects, but we posit that these defects are results of the early cell specification/determination phenotype. These results demonstrate an early role for CDH11 in NC cell formation and future studies will focus on identifying the mechanisms downstream of CDH11 that control cell survival and migration.

**Figure 7.**
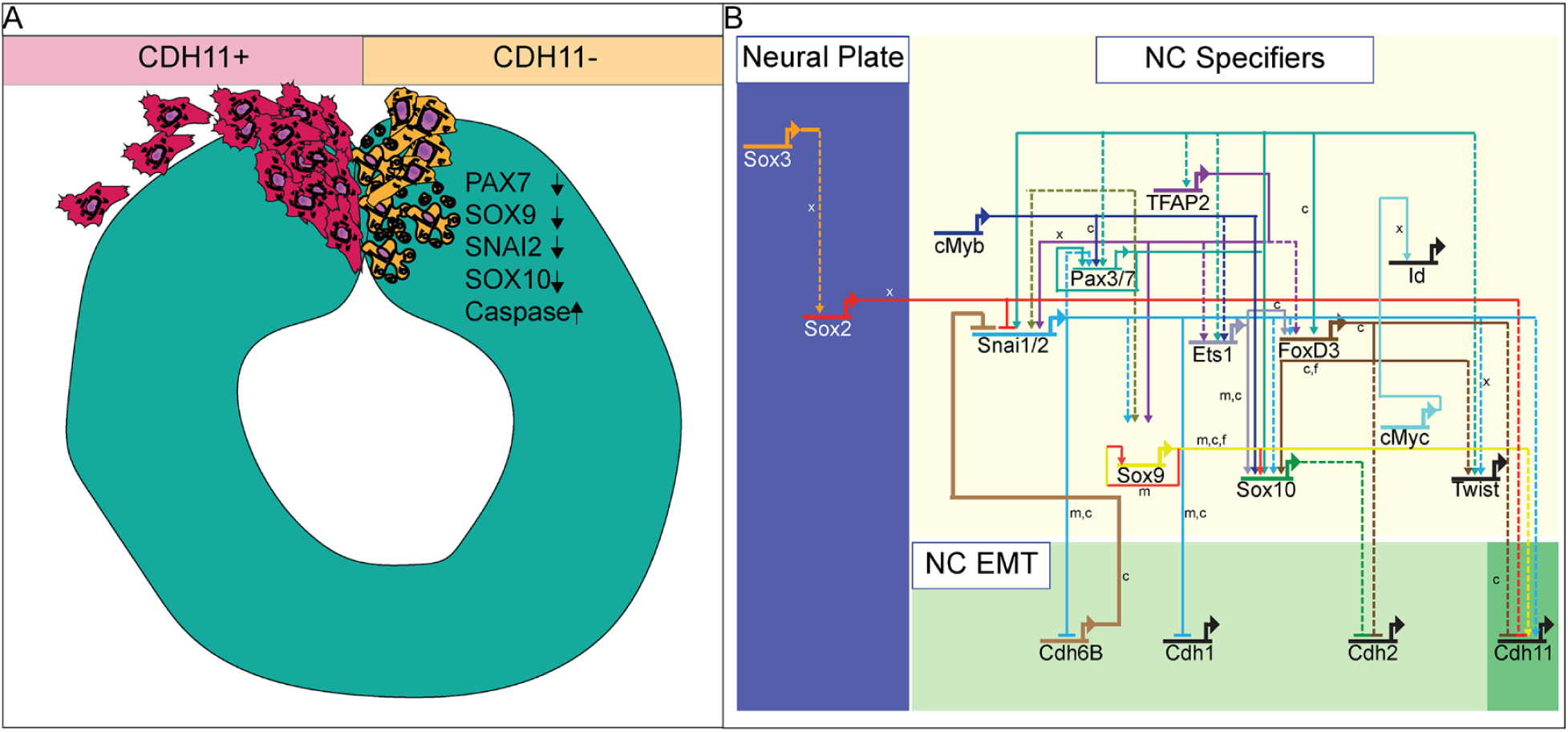
Summary diagram of CDH11-knockdown phenotype. (A) Image depicting NC cell development in normal NC cells (CDH11+) and cells lacking CDH11 (CDH11-). Normal cells undergo EMT, exit the neural tube, migrate collectively, and then progressively mesenchymalize as development proceeds. Normal lamellipodial and filopodial projections form as the cells navigate through the extracellular matrix. In the absence of CDH11, NC cells are induced, but undergo p53-mediated apoptosis due to either 1) inability to complete EMT/migration or 2) altered intracellular signaling in the absence of CDH11. PAX7, SOX9, SNAI2, and SOX10 positive cells are all significantly reduced in the absence of CDH11 while *Casp3 is upregulated. (B) Simplified NC GRN identifying NC specifiers and multiple putative inputs into the CDH11 upstream regulatory region as identified by ATAC-seq performed on premigratory NC cells (Williams et al., 2019). Little is known about the downstream targets and effectors of most cadherin proteins in the NC GRN with the exception of CDH6B (Schiffmacher et al., 2016) and CDH11 in migratory NC cells (Kashef et al., 2009; Koehler et al., 2013), and therefore identifying their targets in premigratory NC cells is essential. Direct binding relationships are indicated with solid lines while putative regulatory relationships are indicated by dashed lines. Letters indicate species in which experiments were performed: x= *Xenopus*, c= chicken, m= mouse. GRN information sourced from (Rogers and Nie, 2018).

### Critical timing of CDH11 function

NC cells are a dynamic population of cells controlled by multiple levels of secreted morphogens, transcription factors, epigenetic modifiers, and cell-cell adhesion molecules (Martik and Bronner, 2017). To understand the potential role of CDH11 in early development, we characterized the spatiotemporal expression and localization of the protein *in vivo* in stages earlier than those previously identified (Chalpe et al., 2010). Our resulting data indicate that CDH11 protein is expressed in the neuroepithelium during neurulation and is specifically upregulated above its neuroepithelial levels in the bonafide NC cells (SOX9+, SNAI2+), rather than all NC progenitors (PAX7+ alone), prior to EMT (Fig. 1). The bulk of previous work studying CDH11 function in NC cells focused on its necessity for normal NC cell migration including its post-translational processing (Abbruzzese et al., 2016; McCusker et al., 2009), the formation of focal adhesions, protrusive activity, and extracellular matrix dynamics (Kashef et al., 2009; Langhe et al., 2016; Row et al., 2016). Here, we focused on understanding the pre-migratory role of CDH11 in NC cell development. CDH11 is absent from the neural plate border cells during NC induction and our results suggest that it is not necessary for NC induction (3 SS and earlier), but rather it is necessary for NC cell specification and/or determination (5 SS and later) (Fig. 2). In support of our results, previous studies in *Xenopus* embryos showed that both exogenous CDH11 and dominant negative CDH11 expression reduced the population of undifferentiated NC cells and inhibited NC cell migration without affecting neural plate specification (Borchers et al., 2001). We believe that our results fill a gap in the previously reported phenotype. Loss of CDH11 reduces both progenitor and definitive NC cell populations (Fig. 2, 3) because without CDH11, these cells undergo p53-mediated cell death (Fig. 4, 5). The timing of cell death occurs after induction but prior to NC cell EMT and migration. Our data suggest one of two possibilities; intracellular signaling downstream of CDH11 is necessary for the maintenance and survival of the premigratory NC cells, or that the cells require the presence of CDH11 for normal cell-cell adhesion-related migration, and without it they die. Future experiments are designed to test those hypotheses. Although CDH11 and other cadherins currently reside at the end of the NC GRN and are thought to function solely as regulators of NC EMT and migration downstream of NC specifiers, recent work that characterized all open enhancers in premigratory NC cells and identified putative binding sites in the CDH11 enhancer for SNAI2 and SOX9 (Fig. 7B) (Williams et al., 2019). Their enhancer analyses suggest that SNAI2 and SOX9 may drive CDH11 expression prior to EMT while FOXD3 and SOX2 may repress it (Williams et al., 2019). Future work will determine which of these factors acts upstream of CDH11 in the NC cells and whether the effects of CDH11 knockdown on cell specification and survival are direct (SOX9/SNAI2→CDH11→cell specification/survival) or indirect through changes in NC specifiers (CDH11→SOX9/SNAI2→cell specification/survival).

### CDH11 and p-53 mediated apoptosis

Our initial investigation into the role of CDH11 during early embryogenesis stemmed from its expression in developing NC cells and the varied accounts of its function in cancer cells. Previous studies in embryos demonstrated that CDH11 is necessary for NC migration (Abbruzzese et al., 2016; Kashef et al., 2009; Langhe et al., 2016; Vallin et al., 1998), but very few studies dissected its function in formation or survival (Borchers et al., 2001). Additionally, the role of CDH11 in cancer cells is variable. In murine retinoblastoma, CDH11 functions as a tumor suppressor. Expression is normally reduced in the retinoblastoma cells, and overexpression of the gene in mouse models resulted in increased cell death (Marchong et al., 2010), however, researchers also noted that in mutant mice lacking CDH11, the number of tumors was reduced. This study supports our data demonstrating that loss of CDH11 induces cell death or reduces NC cell survival, however, the mechanisms driving cell death downstream of gain and loss of CDH11 are likely unique in each situation. Both SLUG/SNAI2 (Tribulo et al., 2004) and SOX9 (Cheung et al., 2005) have demonstrated anti-apoptotic activity in NC cells, and their presence in the neural folds is correlated with reduced cell death. CDH11 knockdown reduces SOX9 and SNAI2 expression significantly (Fig. 3, Fig. 7), which may lead to an indirect activation of the p53-mediated apoptotic pathway.

However, rather than driving cell death indirectly, it is possible that CDH11 is required for normal NC migration, and that loss of CDH11, whether through cell-cell adhesion or intracellular signaling, drives cell death. Loss of CDH11 in the premigratory cells may create a defect in the mechanisms driving NC delamination and EMT. At these stages, the cells have activated NC specifier proteins (SOX9, SNAI2, SOX10) and attempt to migrate, but cannot due to a lack of collective cell migratory ability. Cells lacking CDH11 have increased F-actin cabling (Fig. 6), which has previously been linked to the execution phase of cell death and is required for the formation of apoptotic bodies in embryonic carcinoma cells (Neradil et al., 2005). Additionally, actin-induced activation of the Ras signaling pathway has been previously linked to apoptosis (Gourlay and Ayscough, 2006). In contrast, overexpression of CDH11 may function to activate cell death via a Wnt-or Rho-dependent mechanism as previously described (Li et al., 2012; Row et al., 2016), and future experiments are designed to identify CDH11-specific apoptotic mechanisms.

### Implications and considerations for disease studies

Contrasting conclusions about the role of CDH11 in the disease state may be clarified if the role is assessed in the context of epithelial vs. mesenchymal cells. Loss of CDH11 prior to NC cell migration prevents the cells from leaving the neural tube efficiently, thereby activating the p53-mediated apoptotic pathway. In these cells, CDH11 expression is upregulated on the membrane just prior to EMT, and is maintained in the cells as they collectively migrate, with subsequent down regulation or internalization as the cells travel further from the midline and mesenchymalize (Fig. 1, Fig. S1). The lack of specific expression in NC progenitors, and generalized neural tube expression until just prior to the stage of migration suggests that CDH11 may play another role in the developing neuroepithelium. These data support a dual role for CDH11 during development. The upregulated expression of CDH11 in premigratory NC cells at 5 SS coincides with the wild type expression of SNAI2, SOX9, and SOX10, three proteins that are necessary for NC cell migration. It also coincides with the stage at which N-cadherin is reduced in the dorsal neural tube (Rogers et al., 2018). We believe that the functional type II cadherin complex is required for the completion of EMT, and that loss of CDH11 leads to an increased tension on the cytoskeletal elements of the NC cells as evidenced by the increased CDH1 expression and F-actin localization causing activation of the p53-mediated apoptotic pathway. This hypothesis is supported by previous work demonstrating tightly controlled cadherin proteins in the NC EMT process (Coles et al., 2007; Rogers et al., 2013; Scarpa et al., 2015; Schiffmacher et al., 2014; Schiffmacher et al., 2016). In essence, the cells are programmed to migrate, but because they are unable, they are directed to die. As altering CDH11 affects cell survival, CDH11 may be functioning as it does in cancer cells, as pro-apoptotic stemness modulator that functions via the WNT and Rho pathways (Li et al., 2012), or it may function to allow. Although both gain and loss of CDH11 induce cell death, the mechanisms are likely disparate. Future experiments will focus on understanding the specific mechanisms downstream of CDH11 that regulate the NC cell population.

### Possible mechanisms downstream of CDH11

In *Xenopus*, CDH11 controls filopodia and lamellipodia formation by binding to the guanine nucleotide exchange factor (GEF) proteins. Overexpression of cdc42, Rac1, and RhoA in frog embryos lacking CDH11, rescued cranial NC cell migration (Kashef et al., 2009). GEF1 activates Rac1 and cdc42, and GEF2 activates RhoA, in conjunction factors mediate cytoskeletal structure during axon guidance (Bateman and Van Vactor, 2001). Therefore, there may be alterations in intracellular signaling and cytoskeletal rearrangements after loss of CDH11 leading to their NC phenotype. Future experiments will be designed specifically to determine if the NC cell death phenotype is linked to altered Rac/Rho signaling downstream of CDH11.

CDH11 also interacts with the proteoglycan, Syndecan-4, which interacts with fibronectin and maintains cell-substrate adhesion during cell migration. Loss of Syndecan-4 increased Rac activity, and inhibition of Rac1 by Syndecan-4 regulated the migration of NC cells (Langhe et al., 2016; Matthews et al., 2008). However, the non-canonical Wnt Planer Cell Polarity (PCP) Pathway promotes RhoA activity, which is necessary for NC cell migration, and inhibition of RhoA increased in Rac activation which can induce cell death (Matthews et al., 2008). It is possible that if the NC phenotype caused by loss of CDH11 is not directly related to cell-cell adhesion specific migration defects, rather, CDH11 may be a novel regulator that functions between Rac1 and RhoA, and loss of CDH11 may activate Rac1, preventing the cells from migrating and inducing apoptosis. Future studies will continue to investigate the mechanisms that cause a reduction in the NC cell population in CDH11-deficient cells, but it is clear from our studies and others, that CDH11 plays a complex and important role in NC cell formation and survival prior to its role in migration.

## Materials and methods

### Chicken embryos

Fertilized chicken eggs were obtained from local sources (Sunstate Ranch, CA and the UC Davis Hopkins Avian Facility) and incubated at 37 °C to the desired stages according to the criteria of Hamburger and Hamilton (HH). Use and experiments on embryos was approved by the California State University Northridge IACUC protocol: 1516-012a, c and the UC Davis IACUC protocol #21448.

### Microinjection and Electroporation

Translation blocking antisense fluorescein or biotin-labeled morpholinos to CDH11 (CDH11MO) (5’-TATTTTGTAGGCACAGGAGTATCCA-3’), p53 (5’-CAATGGTTCCATCTCCTCCGCCATG-3’) and a non-specific control morpholino (ContMO) (5′-CCTCTTACCTCAGTTACAATTTATA-3′) were microinjected into the right side of a Hamburger-Hamilton stage 4-5 chicken embryo and platinum electrodes were placed vertically across the chick embryos and electroporated with five pulses of 6.3–6.8 V in 50 ms at 100 ms intervals. Injections of the morpholinos (0.5 mM–1mM) were paired with 0.5–1.5 mg/ml of carrier plasmid DNA (Voiculescu et al., 2008) to enhance cell uptake of treatment. Injections were performed by air pressure using a glass micropipette targeted to the presumptive neural plate and neural plate border region. DNA plasmids pCAGGS-CDH11-IRES-GFP (VectorBuilder.com), Sirius-H2B-C-10 (injected as marker for CDH11MO) was a gift from Michael Davidson to Addgene (Addgene plasmid # 55226; http://n2t.net/addgene:55226; RRID:Addgene_55226), and were introduced in a similar manner to morpholinos described above. HH stage 4–5 electroporations were conducted on whole chick embryo explants placed ventral side up on filter paper rings.

### Immunohistochemistry

For immunohistochemistry (IHC), chicken embryos were fixed on filter paper in 4% paraformaldehyde (PFA) in phosphate buffer for 15-25 mins at room temperature. After fixation, embryos were washed in TBST + Ca2+ with 0.5% Triton X-100. For blocking, embryos were incubated in TBST + Ca2+ with 0.5% Triton X-100 and 10% donkey serum for at least 1 hr at room temperature. Primary antibodies were diluted in blocking solution and incubated with embryos for 3 hr at room temperature or for 24-48 hr at 4° Celsius. After incubation with primary antibodies, whole embryos were washed in TBST + Ca2+, incubated with AlexaFluor secondary antibodies diluted in blocking buffer for 3 hr at room temperature or 12 hr at 4° C, washed in TBST + Ca2+, and post-fixed in 4% for 30 min-1 hr in PFA at room temperature. Antibodies used in the study: Rabbit α-Cadherin-11 (Cell Signaling Technologies, #4442), Mouse α-Cadherin-11 (Invitrogen, #5B2H5), Mouse α-E-cadherin (BD Transduction Laboratories, 61081), Rabbit α-Active Caspase-3 (R&D Systems, #AF835), Mouse α-PAX7 (DSHB), Rabbit α-SOX9 (EMD Millipore, #ab5535), Rabbit α-SOX2 (Abcam, #ab97959), Mouse α-SOX10 (Proteintech, #66786-1-Ig), Rabbit α-SLUG/SNAI2 (Cell Signaling Technology, #9585S), Rabbit α-Phospho histone H3 (R7D Systems, #ab5176). After IHC all embryos were imaged in both whole mount and transverse section (after cryosectioning) using a Zeiss Imager M2 with Apotome capability and Zen optical processing software.

### Western Blot

Embryo lysate was isolated from 10–20 manually dissected stage HH8–10 chicken heads, neural tubes, or axolotl embryos with associated tissues for Western blot analysis. Lysate was isolated using lysis buffer: 50mM Tris-HCL pH 7.4 with 150mM NaCl plus 1.0% NP-40 and EDTA-free protease inhibitor (Roche cOmplete, #11697498001). SDS page was run on precast 8–12% bis-tris gel (Invitrogen, #NP0321BOX) for 3 h at 60 V, gel was transferred to nitrocellulose using the Invitrogen iBlot2 Dry Blotting System. Nitrocellulose membranes were washed in TBST+Calcium with 0.5% Triton X-100, blocked and incubated with primary antibody in TBST+Calcium with 0.5% Triton X-10 with 5.0% milk or BSA, incubated in (5%) milk protein in (TBST+Calcium) with secondary antibody, and visualized using Prometheus ProSignal Femto ECL Reagent (#: 20-302B) and exposed to Prometheus ProSignal ECL Blotting Film, 5 × 7 in. (#: 30-507L).

### Imaging and Fluorescence Quantification

Fluorescence images were taken using Zeiss ImagerM2 with Apotome.2 and Zen software (Karl Zeiss). Fluorescence was quantified using NIH ImageJ by averaging the relative intensity of 1-6 images per embryo. Background was subtracted uniformly across the images using the background subtraction function in NIH ImageJ with a rolling-ball radius of 50.00 pixels before quantitation (Hutchins and Szaro, 2013). Half embryos injected with CDH11MO or ContMO were compared to the uninjected or control side.

### Ex Vivo Neural Tube Explants

For explant assays, embryos were electroporated with 0.75 mM CDH11MO plus Sirius carrier DNA on the right side of the embryo at HH4. Embryos were cultured until HH8 as described previously (Sauka-Spengler and Barembaum, 2008). At HH8, the neural tubes were dissected out of the embryo in Ringer’s solution and subsequently placed in 8-well chamber slides (Millicell EZ SLIDE 8-well glass, sterile, # PEZGS0816) that were coated with 1% fibronectin. The explants were cultured in DMEM with 10% FBS, 2 mM l-glutamine, and 100 units of penicillin with 0.1 mg/ml streptomycin at 37°C with 0.5% CO2 for 8 h. After incubation, explants were fixed using 4% PFA, washed in TBST + Ca2+, and incubated with phalloidin stain. Cytoskeletal stain was: Invitrogen Molecular Probes Alexa Fluor 568 Phalloidin (#A12380).

### Cell counts and statistical analysis

All experiments were repeated 3-4 times. Cell counts and fluorescence intensity represented in box plots were either performed manually in Adobe Photoshop or were performed using NIH ImageJ. Cell counts were averaged from 1-3 sections per embryo. Mean, median, and standard deviation were calculated using Microsoft Excel across all biological replicates. P-value was calculated in Microsoft excel using a Student’s T-Test with a 2-tailed distribution with unequal variance between samples for stringency. P-values under 0.05 are considered statistically significant.

## Acknowledgements

This research was supported by a National Institute of Health, NICHD grant to CDR (R15HD092170-01) and startup funds from UC Davis. We thank our colleagues from the Rogers Lab at California State University Northridge Department of Biology and those at UC Davis in the Department of Anatomy, Physiology, and Cell Biology Department who provided insight and expertise that greatly assisted the research. The authors declare no competing financial interests.

## Author Contributions

Conceptualization: CDR and AC. Data Curation: AC (functional experiments and imaging), SM (characterization, functional experiments and imaging), CDR (functional experiments, data analysis, and imaging). Formal Analysis (cell counts, statistics): AC, SM, CDR. Funding Acquisition: CDR (funds were provided either by NIH R15 HD092170-01 from the NICHD, CSUN startup funding, and UC Davis startup funding). Methodology: AC, SM, CDR. Supervision: CDR. Writing: CDR (wrote manuscript and performed editing), SM (wrote manuscript and performed editing), AC (performed editing).

