## Supplemental figures and tables for "Cadherin-11 is required for neural crest determination and survival"

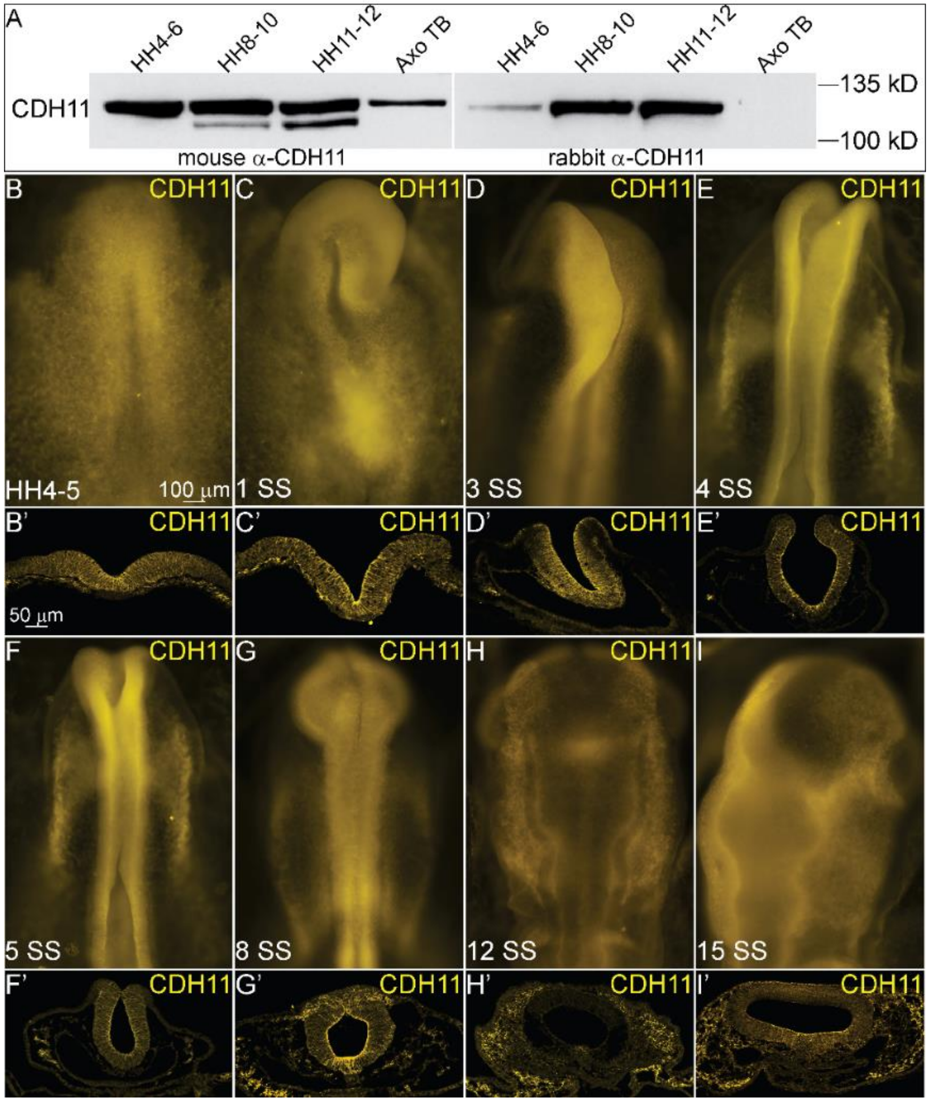

**Supp. Fig. 1. CHD11 expression in whole mount and sections.** To identify spatiotemporal localization in whole mount and in transverse sections, western blot analysis and IHC for CDH11 in protein lysate and uninjected embryos was performed. (A) Western blot analysis using protein lysate isolated from 10 pooled chick or axolotl embryos from stages: HH4-6, HH8-10, HH11-12, and tailbud axolotl embryos. Two antibodies were tested, mouse anti-CDH11 IgG1 or rabbit anti-CDH11 IgG. (B-I) Whole mount and (B'-I' transverse sections of (B-I) after IHC for CDH11 in (B, B') an HH4-5 embryo, (C, C') a 1 SS embryo, (D, D') a 3 SS embryo, (E, E') a 4 SS embryo, (F, F') a 5 SS embryo, (G, G') a 6 SS embryo, (H, H') a 12 SS embryo and (I, I') a 15 SS embryo. Scale bars are as marked (100 µm for whole mount and 50 µm for sections).

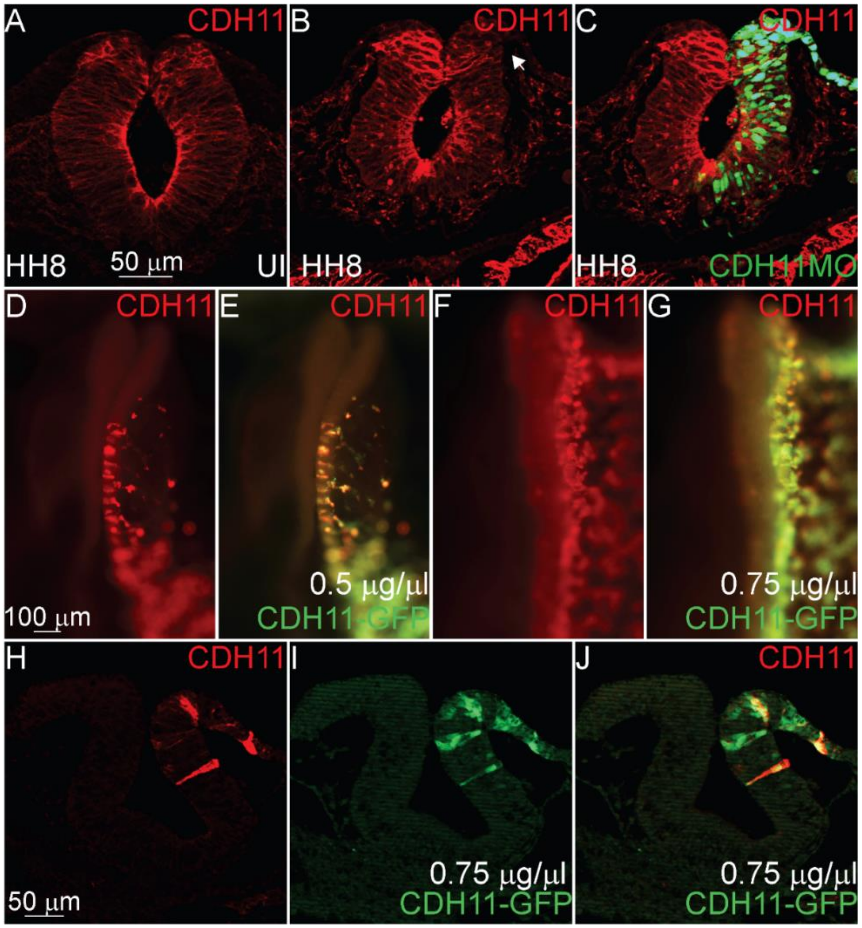

**Supp. Fig. 2. Efficiency of knockdown and overexpression tools.** To verify the efficiency and functionality of CDH11 translation blocking MO and CDH11-GFP full length expression vector, embryos were injected with either treatment and IHC was performed for CHD11. (A) IHC was performed for CDH11 in an uninjected embryo for comparison. (B, C) IHC for CDH11 after CDH11MO electroporation at stage 4 shows that the CDH11MO efficiently inhibits the translation of CDH11 in the injected region of the neural tube. CDH11-GFP was electroporated into stage 4 embryos at 0.5 mg/ml and 0.75 mg/ml, and (D-G) IHC using for CDH11 shows that the CDH11-GFP ectopically expresses CDH11 in the neural tube. (H-J) Section of (F, G) after 0.75 mg/ml CDH11-GFP overexpression. Scale bars are as marked (100 µm for whole mount and 50 µm for sections).

**Supplemental Table 1. Changes in number of PAX7-expressing cells at different stages.**

| HH5 | | | | HH8- | | | | HH8 | | | | HH9+ | | | | Control | | | |
|---|---|---|---|---|---|---|---|---|---|---|---|---|---|---|---|---|---|---|---|
| UI | Ave. # cells per 2-3 serial sections | CDH11MO | Ave. # cells per 2-3 serial sections | UI | Ave. # cells per 2-3 serial sections | CDH11MO | Ave. # cells per 2-3 serial sections | UI | Ave. # cells per 2-3 serial sections | CDH11MO | Ave. # cells per 2-3 serial sections | UI | Ave. # cells per 2-3 serial sections | CDH11MO | Ave. # cells per 2-3 serial sections | UI | Ave. # cells per 2-3 serial sections | ContMO | Ave. # cells per 2-3 serial sections |
| 1 | 27 | 1 | 29 | 1 | 17 | 1 | 19 | 1 | 50 | 1 | 19 | 1 | 58 | 1 | 37 | | 32 | | 34 |
| 2 | 22 | 2 | 18 | 2 | 10 | 2 | 3 | 2 | 54 | 2 | 9 | 2 | 54 | 2 | 32 | | 34 | | 35 |
| 3 | 16 | 3 | 16 | 3 | 23 | 3 | 3 | 3 | 55 | 3 | 43 | 3 | 51 | 3 | 29 | | 34 | | 33 |
| 4 | 23 | 4 | 22 | 4 | 23 | 4 | 11 | 4 | 54 | 4 | 43 | 4 | 64 | 4 | 36 | | 34 | | 23 |
| 5 | 15 | 5 | 14 | 5 | 23 | 5 | 16 | 5 | 59 | 5 | 44 | 5 | 57 | 5 | 39 | | 56 | | 54 |
| 6 | 20 | 6 | 11 | 6 | 21 | 6 | 18 | 6 | 38 | 6 | 17 | 6 | 71 | 6 | 62 | | 56 | | 47 |
| | | | | 7 | 21 | 7 | 13 | 7 | 88 | 7 | 43 | 7 | 37 | 7 | 21 | | 50 | | 49 |
| | | | | 8 | 24 | 8 | 15 | 8 | 83 | 8 | 39 | | | | | | 32 | | 23 |
| | | | | 9 | 16 | 9 | 20 | 9 | 43 | 9 | 22 | | | | | | 14 | | 15 |
| | | | | 10 | 20 | 10 | 15 | 10 | 55 | 10 | 32 | | | | | | 25 | | 20 |
| | | | | 11 | 20 | 11 | 40 | 11 | 31 | 11 | 10 | | | | | | 23 | | 25 |
| | | | | 12 | 28 | 12 | 34 | 12 | 32 | 12 | 17 | | | | | | 33 | | 41 |
| | | | | 13 | 28 | 13 | 48 | 13 | 29 | 13 | 22 | | | | | | 36 | | 41 |
| | | | | 14 | 26 | 14 | 41 | 14 | 29 | 14 | 21 | | | | | | 43 | | 41 |
| | | | | | | | | 15 | 54 | 15 | 32 | | | | | | | | |
| | | | | | | | | 16 | 51 | 16 | 29 | | | | | | | | |
| | | | | | | | | 17 | 64 | 17 | 36 | | | | | | | | |
| | | | | | | | | 18 | 57 | 18 | 39 | | | | | | | | |
| | | | | | | | | 19 | 15 | 19 | 51 | | | | | | | | |
| Mean | 20.50 | | 18.33 | Mean | 21.43 | | 21.14 | Mean | 49.53 | | 29.89 | Mean | 56.00 | | 36.57 | Mean | 35.86 | | 34.36 |
| Median | 21.00 | | 17.00 | Median | 22.00 | | 17.00 | Median | 54.00 | | 32.00 | Median | 57.00 | | 36.00 | Median | 34.00 | | 34.50 |
| Standard | 4.51 | | 6.41 | Standard | 4.85 | | 14.09 | Standard | 18.16 | | 12.60 | Standard | 10.68 | | 12.74 | Standard | 12.00 | | 11.84 |

| | | | | | | | | | | | | | | | | | |
|---|---|---|---|---|---|---|---|---|---|---|---|---|---|---|---|---|---|
| Deviation | | | | Deviation | | | | Deviation | | | | Deviation | | | | Deviation | |
| Student's T-Test (2 tails, type 3) | 0.52 | | | Student's T-Test (2 tails, type 3) | 0.94 | | | Student's T-Test (2 tails, type 3) | 0.0005 | | | Student's T-Test (2 tails, type 3) | 0.01 | | | Student's T-Test (2 tails, type 3) | 0.74 |

**Supplemental Table 2. Changes in number of PAX7, SOX9, SNAI2, and SOX10-expressing cells.**

| PAX7 | | | | PAX7 | | | | SOX9 | | | | SNAI2 | | | | SOX10 | | | |
|---|---|---|---|---|---|---|---|---|---|---|---|---|---|---|---|---|---|---|---|
| UI | Ave. # cells per 2-3 serial sections | CDH11MO | Ave. # cells per 2-3 serial sections | UI | Ave. # cells per 2-3 serial sections | ContMO | Ave. # cells per 2-3 serial sections | UI | Ave. # cells per 2-3 serial sections | CDH11MO | Ave. # cells per 2-3 serial sections | UI | Ave. # cells per 2-3 serial sections | CDH11MO | Ave. # cells per 2-3 serial sections | UI | Ave. # cells per 2-3 serial sections | CDH11MO | Ave. # cells per 2-3 serial sections |
| 1 | 40.33 | 1 | 15.67 | 1 | 32.00 | 1 | 34.00 | 1 | 39.00 | 1 | 22.00 | 1 | 24.00 | 1 | 15.00 | 1 | 23.00 | 1 | 19.00 |
| 2 | 51.00 | 2 | 48.33 | 2 | 34.00 | 2 | 35.00 | 2 | 21.00 | 2 | 9.00 | 2 | 22.00 | 2 | 5.00 | 2 | 26.00 | 2 | 14.00 |
| 3 | 38.00 | 3 | 17.00 | 3 | 34.00 | 3 | 33.00 | 3 | 17.00 | 3 | 6.50 | 3 | 27.00 | 3 | 14.00 | 3 | 27.00 | 3 | 14.00 |
| 4 | 16.50 | 4 | 3.00 | 4 | 34.00 | 4 | 23.00 | 4 | 41.67 | 4 | 19.00 | 4 | 25.00 | 4 | 14.00 | 4 | 22.00 | 4 | 10.00 |
| 5 | 47.00 | 5 | 3.00 | 5 | 56.00 | 5 | 54.00 | 5 | 7.50 | 5 | 7.00 | 5 | 26.00 | 5 | 12.00 | 5 | 16.00 | 5 | 17.00 |
| 6 | 40.33 | 6 | 21.67 | 6 | 56.00 | 6 | 47.00 | 6 | 15.25 | 6 | 12.50 | 6 | 17.00 | 6 | 8.00 | 6 | 23.00 | 6 | 16.00 |
| 7 | 28.67 | 7 | 14.33 | 7 | 50.00 | 7 | 49.00 | 7 | 12.83 | 7 | 8.50 | 7 | 10.00 | 7 | 14.00 | 7 | 11.00 | 7 | 2.00 |
| 8 | 22.00 | 8 | 15.33 | 8 | 32.00 | 8 | 23.00 | 8 | 9.50 | 8 | 5.50 | 8 | 17.00 | 8 | 17.00 | 8 | 9.00 | 8 | 1.00 |
| 9 | 85.50 | 9 | 41.00 | 9 | 14.00 | 9 | 15.00 | 9 | 14.00 | 9 | 12.33 | 9 | 23.00 | 9 | 9.00 | 9 | 9.00 | 9 | 1.00 |
| 10 | 24.67 | 10 | 21.00 | 10 | 25.00 | 10 | 20.00 | 10 | 16.50 | 10 | 12.50 | 10 | 16.00 | 10 | 7.00 | 10 | 6.00 | 10 | 3.00 |
| 11 | 56.50 | 11 | 34.00 | 11 | 23.00 | 11 | 25.00 | 11 | 29.50 | 11 | 15.50 | 11 | 13.00 | 11 | 6.00 | 11 | 6.00 | 11 | 3.00 |
| 12 | 71.00 | 12 | 62.00 | 12 | 33.00 | 12 | 41.00 | | | | | 12 | 29.00 | 12 | 9.00 | 12 | 9.67 | 12 | 4.58 |
| 13 | 37.00 | 13 | 21.00 | 13 | 36.00 | 13 | 41.00 | | | | | 13 | 25.00 | 13 | 18.00 | 13 | 27.00 | 13 | 9.00 |
| 14 | 41.50 | 14 | 25.00 | 14 | 43.00 | 14 | 41.00 | | | | | 14 | 42.00 | 14 | 31.00 | 14 | 20.00 | 14 | 10.00 |
| 15 | 64.00 | 15 | 49.50 | | | | | | | | | 15 | 29.00 | 15 | 21.00 | 15 | 18.00 | 15 | 17.00 |
| 16 | 27.00 | 16 | 44.50 | | | | | | | | | 16 | 19.00 | 16 | 20.00 | 16 | 17.00 | 16 | 11.00 |
| 17 | 21.00 | 17 | 41.67 | | | | | | | | | | | | | 17 | 37.00 | 17 | 19.00 |
| 18 | 58.00 | 18 | 37.00 | | | | | | | | | | | | | 18 | 30.00 | 18 | 26.00 |
| | | | | | | | | | | | | | | | | 19 | 50.00 | 19 | 29.00 |

| | | | | | | | | | | | | | | | | | | | |
|---|---|---|---|---|---|---|---|---|---|---|---|---|---|---|---|---|---|---|---|
| Mean | 42.78 | | 28.61 | Mean | 35.86 | | 34.36 | Mean | 20.34 | | 11.85 | Mean | 22.75 | | 13.75 | Mean | 20.35 | | 11.87 |
| Median | 40.33 | | 23.33 | Median | 34.00 | | 34.50 | Median | 16.50 | | 12.33 | Median | 23.50 | | 14.00 | Median | 20.00 | | 11.00 |
| Standard | 18.82 | | 16.78 | Standard | 12.00 | | 11.84 | Standard Deviation | 11.48 | | 5.29 | Standard Deviation | 7.63 | | 6.75 | Standard Deviation | 11.29 | | 8.28 |
| Student's | 0.02 | | | Student's | 0.74 | | | Student's T-Test (2 | 0.04 | | | Student's T-Test (2 | 0.001 | | | Student's T-Test (2 | 0.01 | | |

**Supplemental Table 3. Changes in number of Caspase-expressing apoptotic bodies and PH3-cells.**

| CASPASE | | | | CASPASE | | | | PH3 | | | | PH3 | | | |
|---|---|---|---|---|---|---|---|---|---|---|---|---|---|---|---|
| UI | Ave. # apoptotic bodies per 2-3 serial sections | Cad11MO | Ave. # apoptotic bodies per 2-3 serial sections | UI | Ave. # apoptotic bodies per 2-3 serial sections | ContMO | Ave. # apoptotic bodies per 2-3 serial sections | UI | Ave. # cells per 2-3 serial sections | Cad11MO | Ave. # cells per 2-3 serial sections | UI | Ave. # cells per 2-3 serial sections | ContMO | Ave. # cells per 2-3 serial sections |
| 1 | 2.00 | 1 | 46.00 | 1 | 13.00 | 1 | 17.00 | 1 | 1.67 | 1 | 5.00 | 1 | 5.00 | 1 | 4.00 |
| 2 | 7.00 | 2 | 28.00 | 2 | 4.00 | 2 | 8.00 | 2 | 4.00 | 2 | 3.75 | 2 | 15.00 | 2 | 7.00 |
| 3 | 37.50 | 3 | 31.00 | 3 | 2.00 | 3 | 6.00 | 3 | 5.00 | 3 | 2.50 | 3 | 3.00 | 3 | 3.00 |

| | | | | | | | | | | | | | | | |
|---|---|---|---|---|---|---|---|---|---|---|---|---|---|---|---|
| 4 | 18.67 | 4 | 29.67 | 4 | 4.00 | 4 | 6.00 | 4 | 8.67 | 4 | 3.67 | 4 | 7.00 | 4 | 6.00 |
| 5 | 14.67 | 5 | 27.33 | 5 | 5.00 | 5 | 10.00 | 5 | 3.33 | 5 | 2.00 | 5 | 3.00 | 5 | 1.00 |
| 6 | 4.67 | 6 | 8.00 | 6 | 5.00 | 6 | 8.00 | 6 | 1.67 | 6 | 1.33 | 6 | 1.00 | 6 | 2.00 |
| 7 | 2.50 | 7 | 19.50 | 7 | 5.00 | 7 | 9.00 | 7 | 0.50 | 7 | 1.50 | 7 | 6.00 | 7 | 7.00 |
| 8 | 30.00 | 8 | 68.50 | 8 | 10.00 | 8 | 13.00 | 8 | 2.50 | 8 | 2.50 | 8 | 7.00 | 8 | 8.00 |
| 9 | 10.00 | 9 | 16.00 | 9 | 10.00 | 9 | 9.00 | 9 | 0.00 | 9 | 6.00 | | | | |
| 10 | 46.00 | 10 | 52.00 | 10 | 16.00 | 10 | 3.00 | 10 | 8.50 | 10 | 6.00 | | | | |
| 11 | 3.00 | 11 | 15.00 | 11 | 13.00 | 11 | 10.00 | 11 | 17.00 | 11 | 17.00 | | | | |
| 12 | 33.00 | 12 | 62.00 | 12 | 20.00 | 12 | 14.00 | 12 | 0.00 | 12 | 2.00 | | | | |
| 13 | 20.67 | 13 | 37.00 | 13 | 9.00 | 13 | 8.00 | 13 | 2.00 | 13 | 4.00 | | | | |
| 14 | 23.00 | 14 | 6.00 | 14 | 15.00 | 14 | 9.00 | 14 | 7.33 | 14 | 11.33 | | | | |
| | | | | | | | | 15 | 4.00 | 15 | 2.33 | | | | |
| | | | | | | | | 16 | 14.33 | 16 | 21.67 | | | | |
| | | | | | | | | 17 | 9.50 | 17 | 10.50 | | | | |
| | | | | | | | | 18 | 12.00 | 18 | 11.33 | | | | |
| | | | | | | | | 19 | 3.67 | 19 | 5.33 | | | | |
| | | | | | | | | 20 | 0.00 | 20 | 5.00 | | | | |
| Mean | 18.05 | | 31.86 | Mean | 9.36 | | 9.29 | Mean | 5.28 | | 6.24 | Mean | 5.88 | | 4.75 |
| Median | 16.67 | | 28.83 | Median | 9.50 | | 9.00 | Median | 3.83 | | 4.50 | Median | 5.50 | | 5.00 |
| Standard Deviation | 14.31 | | 19.32 | Standard Deviation | 5.44 | | 3.54 | Standard Deviation | 4.97 | | 5.48 | Standard Deviation | 4.26 | | 2.60 |

| | | | | | | | | | | | | | |
|---|---|---|---|---|---|---|---|---|---|---|---|---|---|
| Student's T-Test (2 tails, type 3) | 0.042 | | | Student's T-Test (2 tails, type 3) | 0.97 | | | Student's T-Test (2 tails, type 3) | 0.57 | | | Student's T-Test (2 tails, type 3) | 0.54 |

**Supplemental Table 4. Changes in number of PAX7-expressing cells after rescue.**

| PAX7 | | | | PAX7 | | | | PAX7 | | | |
|---|---|---|---|---|---|---|---|---|---|---|---|
| **UI** | **Ave. # cells per 1-3 serial sections** | **CDH11 MO** | **Ave. # cells per 1-3 serial sections** | **UI** | **Ave. # cells per 2-3 serial sections** | **CDH11MO +CDH11** | **Ave. # cells per 1-3 serial sections** | **UI** | **Ave. # cells per 1-3 serial sections** | **CDH11MO +p53MO** | **Ave. # cells per 1-3 serial sections** |
| 1 | 40.33 | 1 | 15.67 | 1 | 37.00 | 1 | 62.00 | 1 | 54.00 | 1 | 19.00 |
| 2 | 51.00 | 2 | 48.33 | 2 | 41.00 | 2 | 58.00 | 2 | 61.00 | 2 | 6.00 |
| 3 | 38.00 | 3 | 17.00 | 3 | 53.00 | 3 | 64.00 | 3 | 33.00 | 3 | 22.00 |
| 4 | 16.50 | 4 | 3.00 | 4 | 46.00 | 4 | 51.00 | 4 | 38.00 | 4 | 34.00 |
| 5 | 47.00 | 5 | 3.00 | 5 | 45.00 | 5 | 46.00 | 5 | 21.00 | 5 | 40.00 |
| 6 | 40.33 | 6 | 21.67 | 6 | 39.00 | 6 | 34.00 | 6 | 26.00 | 6 | 35.00 |
| 7 | 28.67 | 7 | 14.33 | 7 | 80.00 | 7 | 31.00 | 7 | 47.00 | 7 | 67.00 |
| 8 | 22.00 | 8 | 15.33 | 8 | 50.00 | 8 | 39.00 | 8 | 43.00 | 8 | 65.00 |
| 9 | 85.50 | 9 | 41.00 | 9 | 38.00 | 9 | 50.00 | 9 | 7.00 | 9 | 10.00 |
| 10 | 24.67 | 10 | 21.00 | 10 | 49.00 | 10 | 63.00 | 10 | 4.00 | 10 | 8.00 |
| 11 | 56.50 | 11 | 34.00 | 11 | 63.00 | 11 | 101.00 | 11 | 41.00 | 11 | 40.00 |
| 12 | 71.00 | 12 | 62.00 | 12 | 39.00 | 12 | 62.00 | 12 | 48.00 | 12 | 79.00 |
| 13 | 37.00 | 13 | 21.00 | 13 | 41.00 | 13 | 83.00 | 13 | 31.00 | 13 | 44.00 |
| 14 | 41.50 | 14 | 25.00 | | | | | 14 | 40.00 | 14 | 53.00 |
| 15 | 64.00 | 15 | 49.50 | | | | | 15 | 39.00 | 15 | 66.00 |

| | | | | | | | | | | | |
|---|---|---|---|---|---|---|---|---|---|---|---|
| 16 | 27.00 | 16 | 44.50 | | | | | 16 | 28.00 | 16 | 51.00 |
| 17 | 21.00 | 17 | 41.67 | | | | | | | | |
| 18 | 58.00 | 18 | 37.00 | | | | | | | | |
| Mean | 42.78 | | 28.61 | Mean | 47.77 | | 57.23 | Mean | 35.06 | | 39.94 |
| Median | 40.33 | | 23.33 | Median | 45.00 | | 58.00 | Median | 38.50 | | 40.00 |
| Standard Deviatio | 18.82 | | 16.78 | Standard Deviatio | 12.15 | | 19.33 | Standard Deviatio | 15.47 | | 22.75 |
| Student's T-Test [illegible] | 0.02 | | | Student's T-Test [illegible] | 0.15 | | | Student's T-Test [illegible] | 0.48 | | |

**Supplemental Table 5. Changes in fluorescence and migration distance in vivo and cell size and migration ex vivo.**

| CDH1 | | | | PAX7 | | | | SOX9 | | | | Explant: Cell Size | | | | Explant: Distance Migrated | | | |
|---|---|---|---|---|---|---|---|---|---|---|---|---|---|---|---|---|---|---|---|
| UI | Corrected total cell fluorescence | CDH 11MO | Corrected total cell fluorescence | UI | Distance migrated (µm) | CDH 11MO | Distance migrated (µm) | UI | Distance migrated (µm) | CDH 11MO | Distance migrated (µm) | UI | Cell length (µm) | CDH 11MO | Cell length (µm) | UI | Distance migrated (µm) | CDH 11MO | Distance migrated (µm) |
| 1 | 1425.13 | 1 | 1600.39 | 1 | 144.70 | 1 | 74.95 | 1 | 189.11 | 1 | 103.62 | 1 | 33.03 | 1 | 14.72 | 1 | 86.79 | 1 | 78.82 |
| 2 | 1865.40 | 2 | 2761.33 | 2 | 103.02 | 2 | 50.01 | 2 | 170.39 | 2 | 65.80 | 2 | 22.75 | 2 | 31.56 | 2 | 67.63 | 2 | 85.84 |
| 3 | 1125.86 | 3 | 1274.54 | 3 | 159.30 | 3 | 70.19 | 3 | 121.43 | 3 | 91.39 | 3 | 36.63 | 3 | 21.34 | 3 | 78.59 | 3 | 67.02 |

| | | | | | | | | | | | | | | | | | | | |
|---|---|---|---|---|---|---|---|---|---|---|---|---|---|---|---|---|---|---|---|
| 4 | 1367.55 | 4 | 1488.37 | 4 | 162.43 | 4 | 72.81 | 4 | 36.33 | 4 | 32.01 | 4 | 28.30 | 4 | 23.67 | 4 | 100.18 | 4 | 59.00 |
| 5 | 1846.43 | 5 | 2125.36 | 5 | 86.73 | 5 | 56.88 | 5 | 181.28 | 5 | 168.75 | 5 | 57.43 | 5 | 16.05 | 5 | 66.13 | 5 | 80.04 |
| 6 | 1400.92 | 6 | 2258.14 | 6 | 154.02 | 6 | 119.48 | 6 | 169.58 | 6 | 133.15 | 6 | 59.09 | 6 | 11.19 | 6 | 47.44 | 6 | 48.79 |
| 7 | 1275.91 | 7 | 2657.57 | 7 | 189.67 | 7 | 142.44 | 7 | 152.03 | 7 | 149.30 | 7 | 30.04 | 7 | 17.78 | 7 | 48.78 | 7 | 44.94 |
| 8 | 2157.31 | 8 | 2438.92 | 8 | 188.96 | 8 | 39.27 | 8 | 160.64 | 8 | 136.67 | 8 | 37.75 | 8 | 12.75 | 8 | 55.61 | 8 | 41.47 |
| 9 | 2371.54 | 9 | 3222.17 | 9 | 247.83 | 9 | 206.01 | 9 | 253.09 | 9 | 215.33 | 9 | 25.42 | 9 | 9.22 | 9 | 87.52 | 9 | 43.68 |
| 10 | 2646.83 | 10 | 2648.73 | 10 | 199.09 | 10 | 106.24 | 10 | 102.43 | 10 | 62.28 | 10 | 22.90 | 10 | 11.73 | 10 | 94.68 | 10 | 56.76 |
| 11 | 2067.15 | 11 | 2850.50 | 11 | 142.59 | 11 | 61.82 | 11 | 124.75 | 11 | 73.99 | 11 | 52.54 | 11 | 10.83 | 11 | 86.54 | 11 | 67.78 |
| 12 | 1958.55 | 12 | 2772.24 | | | | | 12 | 187.73 | 12 | 111.10 | 12 | 35.18 | 12 | 12.60 | 12 | 43.16 | 12 | 59.58 |
| 13 | 2147.77 | 13 | 2160.19 | | | | | 13 | 185.59 | 13 | 123.45 | 13 | 23.49 | 13 | 30.26 | 13 | 63.05 | 13 | 58.67 |
| | | | | | | | | 14 | 154.84 | 14 | 110.03 | 14 | 41.85 | 14 | 13.89 | 14 | 94.66 | 14 | 57.61 |
| | | | | | | | | 15 | 185.65 | 15 | 142.77 | 15 | 49.38 | 15 | 17.84 | 15 | 61.42 | 15 | 41.96 |
| | | | | | | | | 16 | 180.16 | 16 | 86.04 | 16 | 36.34 | 16 | 19.07 | 16 | 54.41 | 16 | 42.75 |
| | | | | | | | | 17 | 317.04 | 17 | 232.10 | 17 | 34.79 | 17 | 26.30 | 17 | 116.65 | 17 | 61.65 |
| | | | | | | | | 18 | 176.29 | 18 | 104.15 | 18 | 43.96 | 18 | 13.06 | | | | |
| | | | | | | | | 19 | 122.12 | 19 | 47.26 | 19 | 32.89 | 19 | 40.49 | | | | |
| | | | | | | | | | | | | 20 | 28.61 | 20 | 21.18 | | | | |
| | | | | | | | | | | | | 21 | 34.43 | 21 | 15.61 | | | | |
| | | | | | | | | | | | | 22 | 45.47 | 22 | 20.22 | | | | |

| | | | | | | | | | | | | | | | | | | | |
|---|---|---|---|---|---|---|---|---|---|---|---|---|---|---|---|---|---|---|---|
| | | | | | | | | | | | | 23 | 34.74 | 23 | 26.44 | | | | |
| Mean | 1819.72 | | 2327.57 | Mean | 161.67 | | 90.92 | Mean | 166.87 | | 115.22 | Mean | 36.83 | | 19.03 | Mean | 73.72 | | 58.61 |
| Median | 1865.40 | | 2438.92 | Median | 159.30 | | 72.81 | Median | 170.39 | | 110.03 | Median | 34.79 | | 17.78 | Median | 67.63 | | 58.67 |
| Standard Deviation | 466.03 | | 584.50 | Standard Deviation | 44.86 | | 49.30 | Standard Deviation | 57.52 | | 52.45 | Standard Deviation | 10.47 | | 7.83 | Standard Deviation | 21.33 | | 13.94 |
| Student's T-Test (2 tails, type 3) | 0.02 | | | Student's T-Test (2 tails, type 3) | 0.002 | | | Student's T-Test (2 tails, type 3) | 0.006 | | | Student's T-Test (2 tails, type 3) | 0.0000 | | | Student's T-Test (2 tails, type 3) | 0.02 | | |

**Supplemental Table 6. Changes in number of Caspase-expressing apoptotic bodies after rescue.**

| Caspase | | | | Caspase | | | | Caspase | | | |
|---|---|---|---|---|---|---|---|---|---|---|---|
| UI | Ave. # apoptotic bodies per 1-3 serial sections | CDH11 MO | Ave. # apoptotic bodies per 1-3 serial sections | UI | Ave. # apoptotic bodies per 1-3 serial sections | p53 MO | Ave. # apoptotic bodies per 1-3 serial sections | UI | Ave. # apoptotic bodies per 1-3 serial sections | CDH11MO +p53MO | Ave. # apoptotic bodies per 1-3 serial sections |
| 1 | 0.00 | 1 | 46.00 | 1 | 63.00 | 1 | 56.00 | 1 | 35.00 | 1 | 12.00 |
| 2 | 7.00 | 2 | 28.00 | 2 | 55.00 | 2 | 75.00 | 2 | 5.00 | 2 | 2.00 |
| 3 | 37.50 | 3 | 31.00 | 3 | 23.00 | 3 | 24.00 | 3 | 5.00 | 3 | 3.00 |
| 4 | 18.67 | 4 | 29.67 | 4 | 84.00 | 4 | 36.00 | 4 | 15.00 | 4 | 12.00 |
| 5 | 14.67 | 5 | 27.33 | 5 | 42.00 | 5 | 30.00 | 5 | 19.00 | 5 | 22.00 |
| 6 | 4.67 | 6 | 8.00 | 6 | 32.00 | 6 | 35.00 | 6 | 1.00 | 6 | 5.00 |
| 7 | 2.50 | 7 | 19.50 | 7 | 32.00 | 7 | 48.00 | 7 | 24.00 | 7 | 8.00 |
| 8 | 30.00 | 8 | 68.50 | 8 | 25.00 | 8 | 26.00 | 8 | 47.00 | 8 | 38.00 |
| 9 | 10.00 | 9 | 16.00 | 9 | 35.00 | 9 | 30.00 | 9 | 23.00 | 9 | 35.00 |
| 10 | 46.00 | 10 | 52.00 | 10 | 11.00 | 10 | 8.00 | 10 | 16.00 | 10 | 26.00 |

| | | | | | | | | | | | |
|---|---|---|---|---|---|---|---|---|---|---|---|
| 11 | 3.00 | 11 | 15.00 | | | | | 11 | 2.00 | 11 | 1.00 |
| 12 | 33.00 | 12 | 62.00 | | | | | 12 | 6.00 | 12 | 6.00 |
| 13 | 20.67 | 13 | 37.00 | | | | | | | | |
| 14 | 23.00 | 14 | 6.00 | | | | | | | | |
| Mean | 17.90 | | 31.86 | Mean | 40.20 | | 36.80 | Mean | 16.50 | | 14.17 |
| Median | 16.67 | | 28.83 | Median | 33.50 | | 32.50 | Median | 15.50 | | 10.00 |
| Standard Deviatio | 14.49 | | 19.32 | Standard Deviatio | 21.61 | | 18.74 | Standard Deviatio | 14.16 | | 12.96 |
| Student's T-Test | 0.04 | | | Student's T-Test | 0.71 | | | Student's T-Test | 0.68 | | |
